# Molecular mechanism of co-transcriptional H3K36 methylation by SETD2

**DOI:** 10.1101/2024.12.13.628244

**Authors:** James L. Walshe, Moritz Ochmann, Ute Neef, Olexandr Dybkov, Christian Dienemann, Christiane Oberthür, Aiturgan Zheenbekova, Henning Urlaub, Patrick Cramer

**Author notes:** These authors contributed equally (James L. Walshe, Moritz Ochmann). Lead contact: James L.Walshe.

## Abstract

Tri-methylation of histone H3 at residue lysine-36 (H3K36me3) is a hallmark of actively and recently transcribed genes and contributes to cellular memory and identity. The deposition of H3K36me3 occurs co-transcriptionally when the methyltransferase SETD2 associates with RNA polymerase II (Pol II). Here we present three cryo-EM structures of SETD2 bound to Pol II elongation complexes at different states of nucleosome passage. Together with functional probing, our results suggest a 3-step mechanism of transcription-coupled H3K36me3 deposition. First, binding to the elongation factor SPT6 tethers the catalytic SET domain in proximity to the upstream DNA. Second, Pol II nucleosome passage leads to the transfer of a hexasome from downstream to upstream, poised for methylation. Finally, continued transcription leads to upstream nucleosome reassembly, partial dissociation of the histone chaperone FACT and sequential methylation of both H3 tails, completing H3K36me3 deposition of an upstream nucleosome after Pol II passage.

## INTRODUCTION

During transcription of a gene, RNA polymerase (Pol) II progresses rapidly through regular arrays of nucleosomes. Despite the large-scale movements associated with chromatin transcription, Pol II passage through a nucleosome generally involves transfer of the nucleosome from incoming DNA located in front of Pol II (downstream) to DNA in the wake of Pol II (upstream). Such nucleosome transfer during Pol II passage prevents the loss of information contained in the form of covalent histone modifications^1–5^.

The process of Pol II nucleosome passage requires the concerted action of several elongation factors, chromatin remodelers and histone chaperones^6–10^. Elongation factors associated with activated elongating Pol II include DSIF (SPT4/5), SPT6, TFIIS, and PAF1c (CTR9, PAF1, LEO1, WDR61, CDC73, and RTF1)^10–12^. *In vivo,* SPT6 has the strongest influence on Pol II processivity and has histone chaperone activity^13^. The PAF1c subunit RTF1 has a profound impact on the Pol II elongation rate *in vitro*^7,10,14^ and is implicated in the maintenance of chromatin by recruiting histone remodelers and modifiers^14–21^. In addition to these elongation factors, the histone chaperone FACT (a heterodimer of subunits SPT16 and SSRP1) stimulates chromatin transcription^6,8,22–25^ and is also implicated in chromatin maintenance^26–33^. Recently, Cryo-EM structures of the yeast activated elongation complex transcribing through a nucleosome have progressed our understanding of the mechanisms underlying Pol II chromatin passage^6,27,34,35^.

However, despite this progress, it remains unknown how Pol II nucleosome passage is coupled to co-transcriptional histone modification. One of the most prominent histone modifications acquired during Pol II transcription is the tri-methylation of histone H3 on residue lysine-36 (H3K36me3) that was first described over 20 years ago^36–41^. H3K36me3 acts as docking sites for various chromatin readers, including PWWP, chromo– and tudor domains^42–45^ and is implicated in pre-mRNA processing, DNA mismatch repair^46^ and chromatin integrity^47^. Changes in H3K36me3 levels lead to deleterious intergenic transcription that is associated with cancerous cell growth^48,49^.

The deposition of H3K36me3 is carried out by the conserved histone methyltransferase SETD2^50^. SETD2 associates with Pol II through a interaction of its Set2 RPB1-interaction (SRI) domain with the phosphorylated C-terminal domain (CTD) of the largest Pol II subunit RBP1^39,51,52^. The SRI domain also antagonizes the effect of the SET auto-inhibitory domain (AID)^53^. The yeast homologue of SETD2, Set2, interacts genetically with SPT6^54,55^. To date, structural analysis of SETD2 is limited to individual domains^51,52,56^ and the catalytic SET domain bound to a nucleosome^54^.

In this study, we investigate the molecular-mechanistic basis of co-transcriptional H3K36me3 deposition by SETD2. We demonstrate H3K36me3 deposition occurs in the wake of elongating Pol II, after the nucleosome has been transferred to upstream DNA. We present three cryo-EM structures of the activated Pol II elongation complex in the presence of SETD2 and FACT and probe these structures with functional assays. Our structure-function analysis provides a model for a 3-step molecular mechanism of co-transcriptional H3K36me3 deposition.

## RESULTS

### H3K36me3 deposition requires Pol II nucleosome passage

To investigate the mechanism underlying co-transcriptional H3K36me3 deposition, we first determined whether SETD2 methylates the nucleosome downstream or upstream of the transcribing Pol II. To this end, we performed RNA extension assays on a DNA template wrapped around a single nucleosome positioned 111 bp downstream from the extension start site and monitored H3K36me3 deposition with Western blot analysis. A T-less cassette was used to enable stalling of Pol II at base pair (bp) 32 of a modified Widom 601 sequence (Fig. 1a). Poll II stalled at this position would unwrap DNA to at least superhelical location (SHL) −4, providing an optimal downstream nucleosome substrate for H3K36me3 deposition^57^.

**Figure 1:**
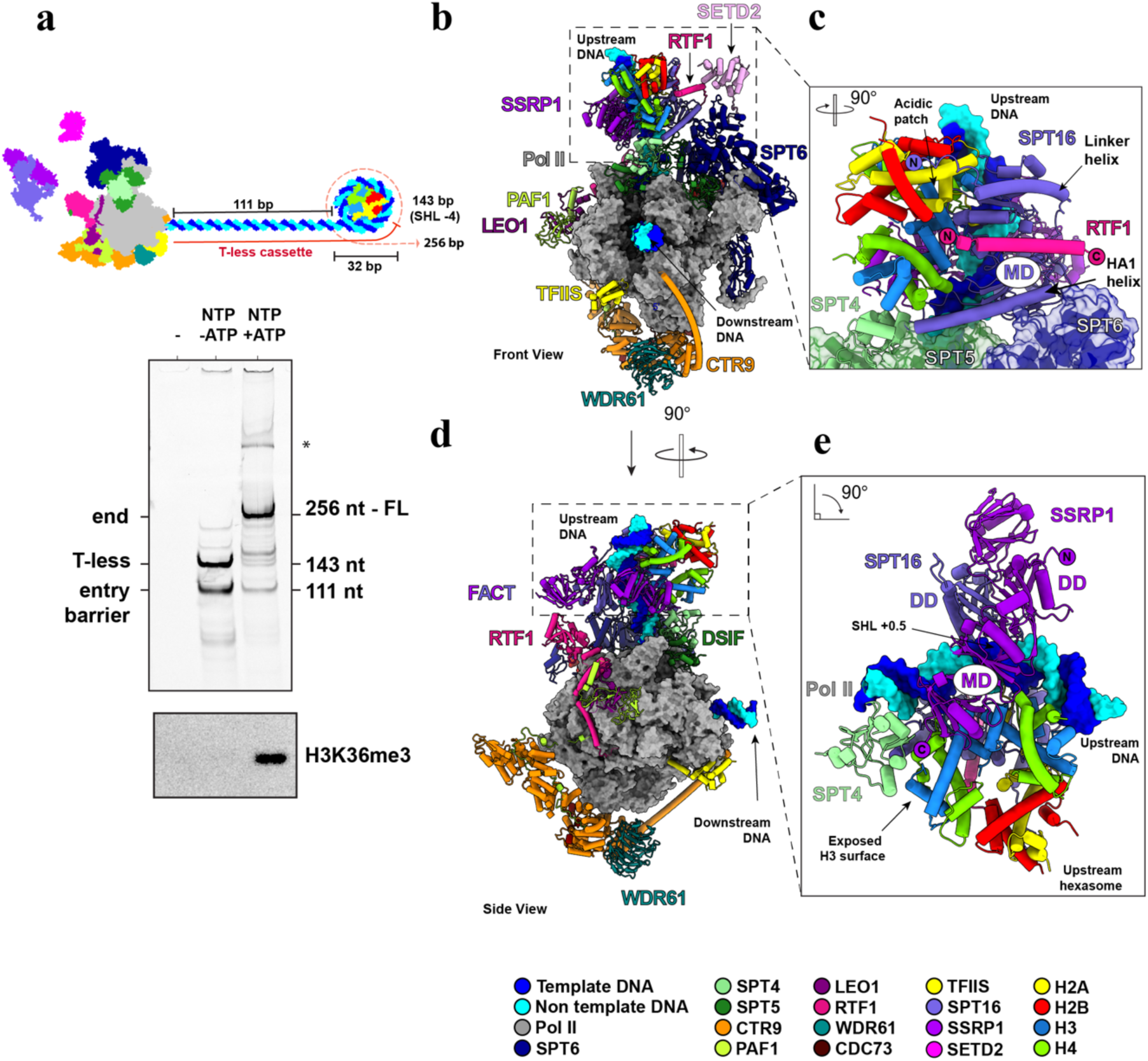
Upstream H3K36me3 activity and Cryo-EM structure the poised methylation complex. **a**, Schematic representation of nucleosome template and proteins used in **b**, SETD2 deposits H3K36me3 on an upstream nucleosome. RNA extension by the methylation competent elongation complex through mononucleosomes with extended run-up distance and T-less cassette to bp 32 (SHL −4) into the nucleosome. RNA primer contains a 5’ Cy5 label. RNA lengths are indicated on the right side of the denaturing PAGE gel. The positions of prominent stalling products are indicated on the left. Western blots of H3K36me3 of RNA extension assy. **c**, Overall structure of the complex. **d**, Magnified view of FACT binding to upstream hexasome. **d**, Alternate view of the complex **e**, RTF1 binding to FACT

The complete mammalian activated elongation complex^10^ was assembled on the nucleosomal DNA construct and the factors TFIIS, FACT, IWS1 and SETD2 (residues 1435-2567) were added (Fig. 1a). H3K36me3 deposition was analyzed 10 minutes after nucleotide addition. No H3K36me3 was detected in the absence of transcription, corroborating that methylation occurs co-transcriptionally in our minimal biochemical system. Also, H3K36me3 was absent after transcription was stalled at bp 143, when Pol II is located in front of the incoming nucleosome. A signal for H3K36me3 was only observed after transcription had occurred to the end of the template (bp 256). This shows that co-transcriptional H3K36me3 deposition occurs on the upstream nucleosome after Pol II passage (Fig. 1a).

### Architecture of the poised methylation complex

In order to investigate how SETD2 methylates an upstream nucleosome, we biochemically and structurally analyzed the progression of the complete mammalian activated elongation complex^10^ in the presence of TFIIS, FACT, IWS1 and SETD2. We stalled the complex at a position that is known to trigger nucleosome transfer upstream of the Pol II (bp 64 of a modified Widom 601 sequence)^27^. The complex was mildly crosslinked and purified in a Grafix gradient. Fractions containing the stalled complex (called here methylation-competent elongation complex) were used for single-particle cryo-EM analysis (Extended Data Fig. 1). We obtained a reconstruction that showed Pol II at a resolution of 2.85 Å (Extended Data Fig. 2). Focused classification of density upstream of Pol II revealed a FACT-bound hexasome and SETD2 bound to SPT6. An overall model for the poised methylation complex (State 1), was built using the obtained density maps (Extended Data Fig. 2-3), data from chemical crosslinking-mass spectrometry (CXL-MS) (Methods) (Fig. 2, Extended Data Fig. 4a-b) and Alphafold2 or ColabFold predictions^58,59^ (Extended Data Fig. 5).

**Figure 2:**
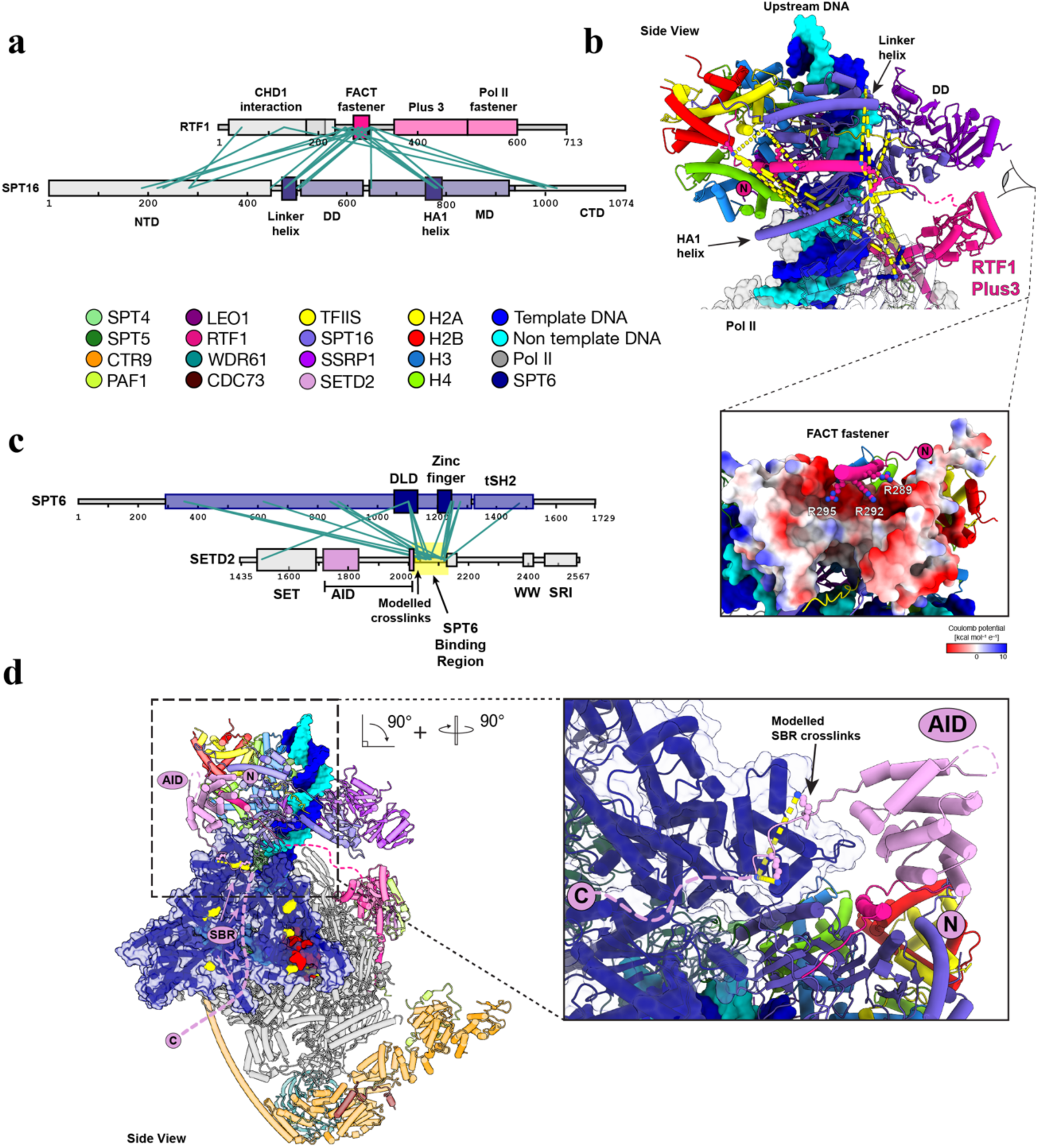
CXL-MS analysis poised methylation complex. **a**, Schematic overview of the RTF1 and SPT16 domain architecture. Obtained BS3 crosslinks are visualized between the proteins and verify interaction between RTF1 FACT “fastener” and SPT16. Only crosslinks with a – log10Score >1 are shown. Crosslinks to the N-terminal remnant of the RTF1 and SPT16 affinity tags are not displayed **b**, Magnified view of FACT “fastener” binding SPT16. Interchain BS3 crosslinks are visualized. SPT16 visualized as Coulombic surface potential. **c**, Schematic overview of the SPT6 and SETD2 domain architecture. Obtained BS3 crosslinks are visualized between the proteins **d**, Overall structure of the complex. BS3 crosslink to SPT6 colored yellow. Magnified view of DLD binding motif of SETD2 AID on surface of SPT6.

In State 1, FACT binds the transferred hexasome in the region of upstream DNA above the Pol II active center cleft (Fig. 1b-e). Relative to the direction of transcription, the hexasome is missing the proximal H2A–H2B dimer and is consistent with a similar state captured for the yeast Pol II^27^. Approximately 20 bp of DNA covers the histones from SHL −0.5 to SHL +1.5 (Fig. 1e). FACT is bound to all histones and the DNA, covering a total of ∼ 5200 Å^2^ in buried surface area. Its dimerization domains (DD) interact with the nucleosomal DNA at SHL + 0.5, approximately 30 bp upstream of the Pol II active center (Fig. 1e). The middle domain (MD) of SPT16, that includes the HA1 helix (residues 759-791), contacts both the (H3–H4)2 tetramer and the distal H2A-H2B dimer (Fig. 1c, Extended Data 4e). The SPT16 linker helix (residues 468-498) extends towards the acidic patch of the H2A–H2B dimer (Fig. 1c) and the residues of the C-terminal domain (CTD) bind the exposed surface of the H2A–H2B dimer (Extended Data Fig. 4d-e) as described^27,60–62^. The SSRP1 middle domain (MD) contacts both the (H3– H4)2 tetramer and DNA on the opposite side to SPT16 (Fig. 1e). Together, these interactions anchor FACT to the hexasome (Fig. 2b-c).

Elongation factors DSIF (SPT4/SPT5) and SPT6 are well positioned to stabilize the FACT-bound upstream hexasome. SPT5 is in close proximity to the SPT16 MD and the HA1 helix (Fig. 1c). SPT4 stabilizes the exposed surface of histone H3 and the C-terminal end of the SSRP1 MD (Fig. 1e). Compared to the canonical complete activated elongation complex^10^, SPT6 rotates towards the upstream DNA by ∼20° (Extended Data Fig. 4f) providing an additional contact point between the elongation complex and the hexasome (Fig. 1c). The absence of this contact coincides with badly resolved nucleosome density (Extended Data Fig. 2) and indicates stabilization of the hexasome by SPT6. This is consistent with the established role for SPT6 as a chaperone^13,63,64^. Based on CXL-MS data (Fig. 2a,b), we placed a previously unresolved helix of RTF1 (residues 266-315) into helical density on the surface of SPT16. This helix is conserved in higher eukaryotes (Extended Data Fig. 6) with basic residues Arg 289, Arg 292 and Arg 295 well positioned to bind to the acidic surface of the middle domain of SPT16 (Fig. 2b, Extended Data Fig. 4c). Due to the direct interaction between RTF1 and FACT, we termed the RTF1 helix the FACT “fastener”. Collectively, we have established in the mammalian context that both SPT6 and FACT fulfill roles as histone chaperones by stabilizing a newly transferred hexasome in the wake of Pol II transcription. Additionally, elongation factors DSIF and RTF1 stabilize the interactions between FACT and the hexasome, demonstrating the intrinsic link between Pol II elongation and nucleosome retention.

### SETD2 binds to SPT6

Careful inspection of our cryo-EM density revealed density in proximity of SPT6 and the upstream hexasome that could not be assigned to a known feature. Further classification on signal-subtracted particles containing only SPT6 showed extra density on the death-like domain (DLD) surface (Extended Data Fig. 7a). This density could be assigned to a conserved 7-residue stretch of SETD2 (Extended Data Fig. 8) based on our CXL-MS data and Colabfold^58^ predictions (Fig. 2c-d, Extended Data Fig. 5b). It is located at the beginning of an unstructured region of SETD2 (residues 2022-2130) that strongly cross-links to SPT6 and was named SPT6 Binding Region (SBR) (Fig. 2c,d). Only 14 residues of this region could be modelled into our observed density, suggesting the remaining unstructured residues are flexible.

The SBR of SETD2 is located C-terminal of the SETD2 auto-inhibitory domain (AID) (Fig. 2c). At low threshold and with a gaussian filter applied, we observed globular density (Extended Data Fig. 7b) into which the Alphafold2^59^ model of the SETD2 AID domain could be fitted. The AID domain is located between the FACT “fastener” of RTF1 and the DLD of SPT6 (Fig. 1b, Fig. 2d, Extended Data Fig. 7b). We did not observe density for the catalytic SET domain, but since it immediately follows the AID (Fig. 2c) the catalytic SET domain is located in proximity to an upstream nucleosome during reassembly. In State 1, the catalytic SET domain is occluded from binding the hexasome by the presence of FACT. In summary, State1 shows the SBR tethers the AID and catalytic SET domain to SPT6 in proximity to the transferred hexasome, poised for H3K36me3 deposition upon FACT dissociation from the dyad.

### Architecture of the proximal H3K36me3 writer complex

Having identified a poised state for H3K36me3 deposition, we investigated how SETD2 would methylate an intact upstream nucleosome. We reconstituted the methylation-competent elongation complex on a ligated nucleosome template that would represent transcription to bp 159 of a Widom 601 sequence, such that the nucleosome is located upstream of Pol II. The complex was purified by size exclusion chromatography and a single peak fraction containing the complex was crosslinked, quenched and flash-cooled for single particle cryo-EM analysis (Extended Data Fig. 9a). The obtained data yielded a reconstruction that revealed Pol II at a resolution of 2.63 Å. Focused 3D classification of the upstream region resulted in a 4.8 Å reconstruction of the catalytic SET domain (residues 1452 – 1696) bound to the nucleosome that we refer to as State 2 (Fig. 3b, Extended Data Fig. 10-11).

**Figure 3:**
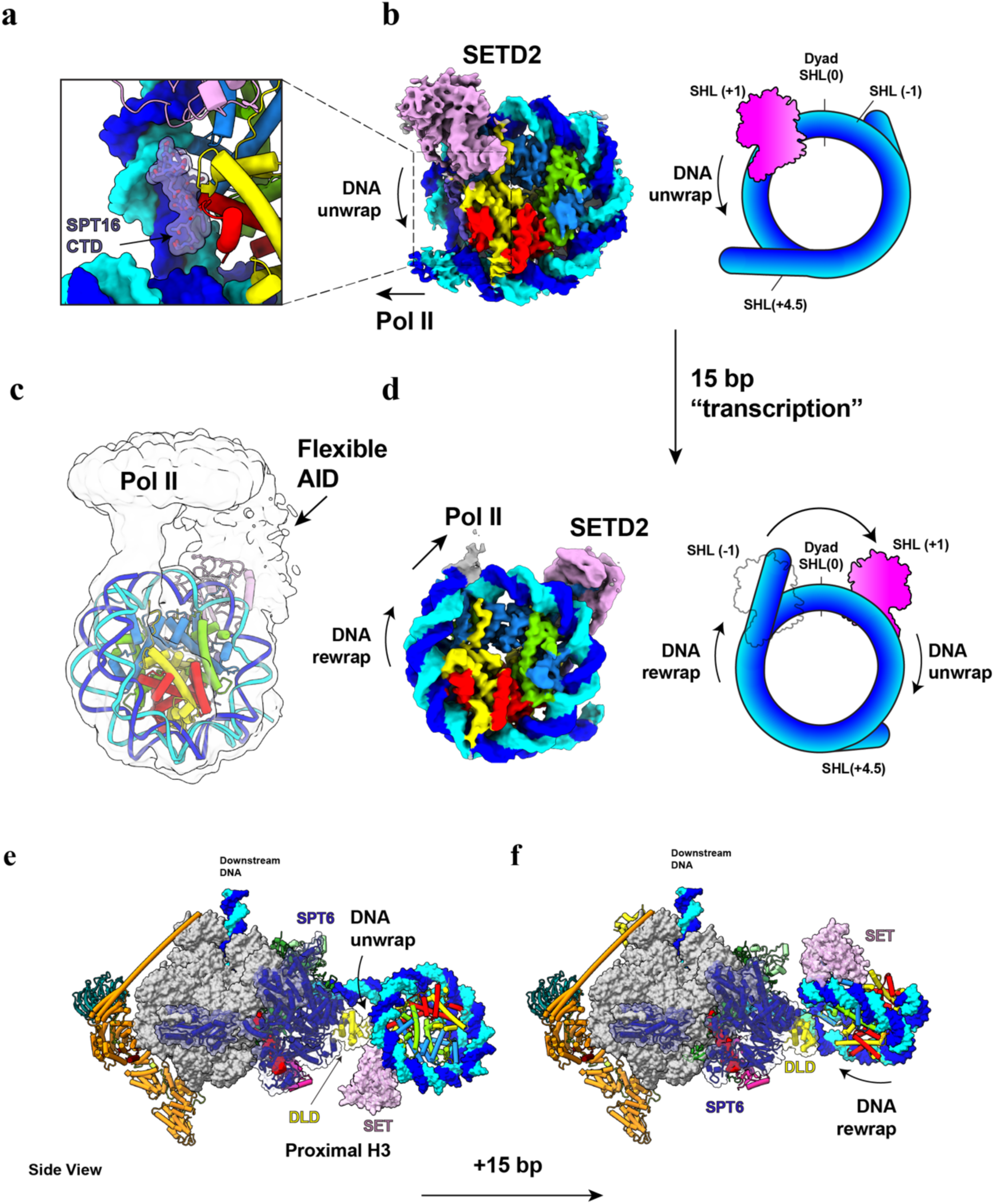
Cryo-EM structure of SETD2 bound to proximal and distal H3 tail. **a**, SPT16 CTD binds exposed H2A–H2B dimer of unwrapped nucleosome. **b**, 4.84 Å reconstruction of SETD2 bound to proximal H3 tail of unwrapped nucleosome **c**, 3.6 Å reconstruction of SETD2 bound to the distal H3 tail of unwrapped nucleosome. A Gaussian filter with a 2σ standard deviation was applied to the cryo-EM map. **d**, As in c, without filter **e,f** Overall architecture of the SETD2 bound to the proximal and distal H3 tails, of an upstream nucleosome, respectively

Compared to the upstream hexasome observed in State 1, in State 2 the catalytic SET domain now binds to the proximal H3 tail at SHL −1 (Fig. 3b). In contrast to previous structures of catalytic SET domains bound to nucleosomes^57,65^, in State 2 the CTD of STP16 (residues 937-951) stabilizes the SETD2-bound partially unwrapped nucleosome (Fig. 3a-b). Although this region is known to bind free H2A–H2B dimers^60,66^, we observe this interaction in the context of a complete nucleosome. This interaction represents a late stage of nucleosome reassembly after FACT deposits the remaining H2A–H2B dimer, prior to complete dissociation. This suggests an additional role for FACT in facilitating H3K36me3 deposition, apart from its known role in stimulating chromatin transcription and maintenance.

Applying a Gaussian filter to the nucleosome reconstruction allowed further tracing of the DNA in one direction, providing an estimate of Pol II direction, and confirming SETD2 is bound to the proximal H3 histone (relative to Pol II) (Extended Data Fig. 12a-b). Surprisingly, adjacent to the catalytic SET domain, above SHL −1, we identified globular density in our cryo-EM map (Extended Data Fig.12b). There are only 20 unstructured amino acids between the last modelled residue of the SET domain and the beginning of the helical AID domain, therefore we assumed this density corresponds to the AID. Consistent with this assumption, the Alphafold2^59^ prediction of the AID fits within the additional density with only 16 Å between the connecting residues (Extended Data Fig. 12a-b). In this conformation, the SBR of SETD2 is directed towards Pol II and SPT6 (Extended Data Fig. 12a-b).

To accurately determine the orientation of the upstream nucleosome relative to Pol II, multibody refinement in RELION^67^ was performed on the subset of Pol II particles that contained well-aligned nucleosomes (Extended Data Fig. 13a). Given the high flexibility, further classification was performed based on the first principal motion competent (Extended Data Fig. 13b). Refinement of selected particles led to a 12.4 Å reconstruction that allowed faithful positioning of both nucleosome and Pol II models (Fig. 3e, Extended Data Fig. 13c). Although the AID remains highly flexible, the C-terminus of the catalytic SET domain is orientated towards the DLD of SPT6 and indicates the AID and SBR of SETD2 likely remain tethered to STP6 (Fig. 3e, Extended Data Fig. 12b-c). Compared to State 1, the overall architecture of State 2 demonstrates that continued transcription would reassemble and position the nucleosome at the upstream edge of Pol II. FACT dissociates from the dyad but remains loosely tethered to stabilize the partially unwrapped nucleosome state. SETD2 is tethered to SPT6 such that the AID does not prevent binding of the catalytic SET domain to the proximal H3 tail of nucleosome.

### Structure of the distal H3K36me3 writer complex

To methylate both H3 tails of an upstream nucleosome, SETD2 would need to access both the proximal and the distal H3 histone tails during transcription. Re-wrapping of the nucleosome observed in State 2 would rotate the nucleosome, with respect to Pol II, such that the distal H3 histone would be in proximity to SPT6. To investigate this, we prepared another complex for cryo-EM analysis on a 15 bp longer ligated nucleosome template, representing transcription to bp 174 into a Widom 601 sequence. Sample preparation, data collection and processing were performed similarly as for the State 2 complex (Extended Data Fig. 9b, 14-15), leading to a reconstruction with a resolution of 2.6 Å for Pol II and 3.6 Å reconstruction of SETD2 bound to the nucleosome.

In the resulting State 3 structure, SETD2 is bound to the distal H3 tail at SHL −1 (Fig. 3c-d). The additional 15 bp of DNA induced approximately 20 bp of DNA to rewrap the nucleosome, making the proximal H3 tail inaccessible for SETD2 binding (Fig. 3c-d). SETD2 binding to the distal H3 tail, at SHL +1, facilitated unwrapping of distal DNA to approximately SHL −4.5. Analysis of the overall architecture of the distal H3K36me3 complex demonstrates the nucleosome rewrapping has rotated the nucleosome by ∼45°, positioning SETD2 on the opposite side of SPT6 (Fig. 3f), such that the interaction between SETD2 and SPT6 could be maintained. Compared to State 2, the overall architecture of State 3 demonstrates that continued transcription would allow rewrapping of the nucleosome that repositions the distal H3 tail of the nucleosome to allow the catalytic SET domain to bind, whilst maintaining an interaction with STP6.

### SETD2-SPT6 interaction is required for H3K36me3 deposition

To test the functional importance of the SETD2 SBR, we designed a series of SETD2 truncations (Fig. 4a) and performed co-transcriptional methylation assays. In these assays we used a chromatinized template consisting of four Widom 601 positioned nucleosomes separated by 30 bp of linker DNA. As a positive control, we included a SETD2 truncation containing only the catalytic SET domain without auto-regulatory elements. As expected, the catalytic SET domain alone (Fig. 4a Construct 1) showed almost three-fold higher methylation levels than WT (Fig. 4b). The addition of the AID abolished methylation (Fig. 4b Construct 2) consistent with its described role in yeast^53^. Consistent with our structural and crosslinking data, a SETD2 variant that additionally includes the SBR (Fig. 4a Construct 3) recovered methylation to approximately ∼50% compared to WT levels. Complete recovery of methylation was not observed as Construct 3 lacks the SRI domain that has been implicated in Set2 recruitment to chromatin ^53,55^.

**Figure 4:**
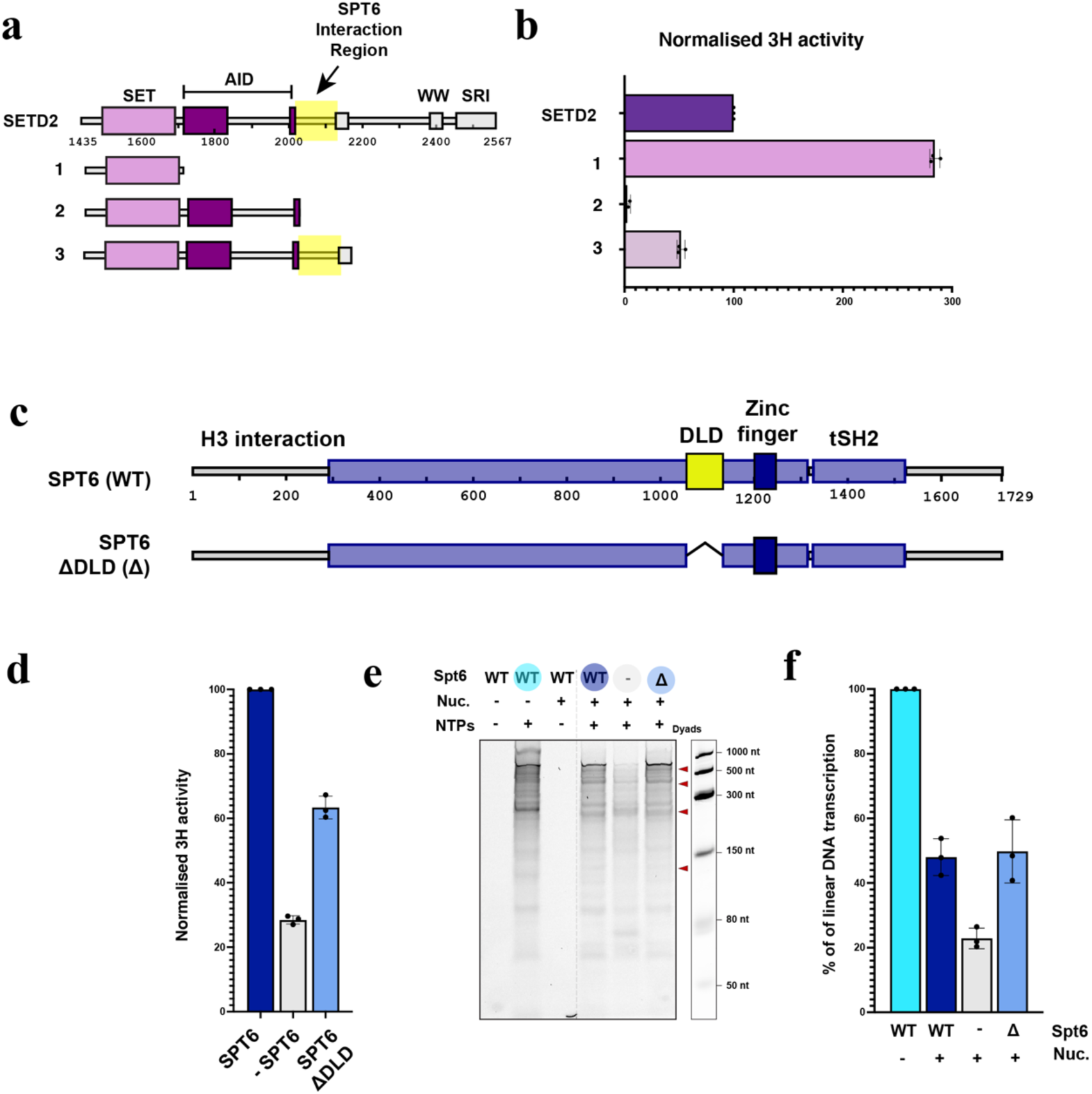
SPT6 SETD2 interaction required for efficient methylation. **a**, Schematic overview of the SETD2 truncations **b**, Normalized 3H activity of co-transcriptional methylation assays performed with SETD2 or truncations. Reactions performed in triplicate. **c**, Schematic overview of the SPT6 showing DLD truncation **d**, Normalized 3H activity of co-transcriptional methylation assays Reactions performed in triplicate. **e**, As in d, with cold S-Adenosyl methionine. Estimate dyad positions indicated on the right. **f**, Quantification of replicate assays in d. For data presented in b,d,c each point reflects one replicate (N=3), depicted as mean ± s.d.

In our structures, we observed 7 residues of the SBR that bind to the DLD of SPT6. We therefore also tested the functional importance of the DLD domain. We generated an SPT6 ΔDLD construct and again performed co-transcriptional methylation assays (Fig. 4c). Omitting SPT6 from the reaction resulted in a ∼75% decrease in methylation activity, whereas the deletion of the DLD resulted in a ∼40% decrease (Fig. 4d). Although these results suggest a role for the DLD in SETD2-dependent histone H3 methylation, SPT6 is an elongation factor and may indirectly facilitate H3K36me3 deposition by increasing transcription that exposes the SETD2 binding sites at SHL+/–1. To test this possibility, transcription activity of elongation complexes formed with full-length SPT6 and ΔDLD mutant were analyzed by denaturing PAGE. The decrease in methylation in the absence of full length SPT6 is likely due, in part, to decreased transcription (Fig. 4e-f), however no change in transcription was observed for the SPT6 ΔDLD, indicating the methylation deficit was caused by the absence of the DLD. These results disentangle SPT6-stimulated transcription from co-transcriptional H3K36me3 deposition, and conclusively show that the SPT6 DLD is required for efficient co-transcriptional H3K36me3 deposition, and strongly support the model that binding of SETD2 to SPT6 via the SBR-DLD interaction is important for high-efficiency H3K36me3 deposition.

## Discussion

To enable co-transcriptional H3K36me3 deposition while preventing spurious methylation events, SETD2 activity is tightly regulated in cells^53,68,69^. In the absence of transcription, the inhibitory effect of the SETD2 AID prevents spurious H3K36me3 deposition on unwrapped nucleosomes that are generated by DNA replication or repair^53^. SETD2 recruitment to chromatin involves an interaction between the SETD2 SRI and the Pol II CTD^36^, but the molecular mechanism to overcome auto-inhibition remained unclear. Here we provide three cryo-EM structures and complementary biochemical data that provide answers to these open questions and expand our understanding of how SETD2 is bound to an activated Pol II elongation complex and how it is positioned for co-transcriptional H3K36me3 deposition.

With respect to binding of SETD2 to the transcribing polymerase complex, we have identified a novel SPT6 Binding Region (SBR) within SETD2 that positions the catalytic SET domain at the upstream edge of Pol II. The direct SPT6 interaction explains the genetic interactions previously observed^54^ and the incomplete depletion of Set2 from chromatin when the SRI domain is removed^55^. In yeast, when only the catalytic SET domain was fused to the CTD of RBP1, no H3K36me3 was detected and indicates the SBR is critical for correct positioning of the catalytic SET domain at the upstream edge of Pol II. When tethered to SPT6, the AID of SETD2 is unable to inhibit binding of the catalytic SET domain to a nucleosome that reassembles in the wake of Pol II transcription. In summary, this structure-function analysis of SETD2 shows both the SRI domain and the newly identified SBR are critical for regulating co-transcription H3K36me3 deposition. In light of these results, our data suggests a simple mechanism that physically couples SETD2 binding to the activated Pol II elongation complex to its functional activation, providing the molecular rational for H3K36me3 deposition to only occur co-transcriptionally.

Our data converge with published results and lead to a three-step model for co-transcriptional H3K36me3 deposition. In the first step (Fig. 5), SETD2 binds to SPT6, through the SBR, and positions the catalytic SET domain to the upstream edge of Pol II. In the second step, transcription of Pol II through the nucleosome goes along with FACT-mediated transfer of the incoming nucleosome from downstream to upstream DNA. This transfer positions a hexasome adjacent to the AID of SETD2. Compared to a similar state observed in yeast ^27^, the FACT-bound hexasome forms additional contacts with the histone chaperone SPT6 and with a previously unresolved RTF1 region we call the FACT “fastener”. The FACT “fastener” likely helps retain FACT-bound nucleosomes in proximity to Pol II during transcription, contributing unexpectedly to a previously suggested mechanism^27^. SPT6 extends the nucleosome cradle described previously^27^, indirectly acting as a histone chaperone. SETD2 is now poised for methylation but FACT occludes the catalytic SET domain from binding. In a third step, further transcription then allows FACT to deposit the missing H2A–H2B dimer and to stabilize the partially unwrapped nucleosome, tethering to SPT6 limits the AID domain allowing the catalytic SET domain to bind and methylate the proximal H3 tail (Fig. 5, State 2). Further transcription enables rewrapping of DNA around the proximal histones and repositioning of SETD2 to the distal H3 tail, facilitating methylation of the distal H3 tail and completing H3K36me3 deposition on the transcribed nucleosome (Fig. 5, State 3). This cycle continues throughout the gene body at each Pol II nucleosome passage event, thus resulting in H3K36me3 deposition in actively transcribed genes.

**Figure 5:**
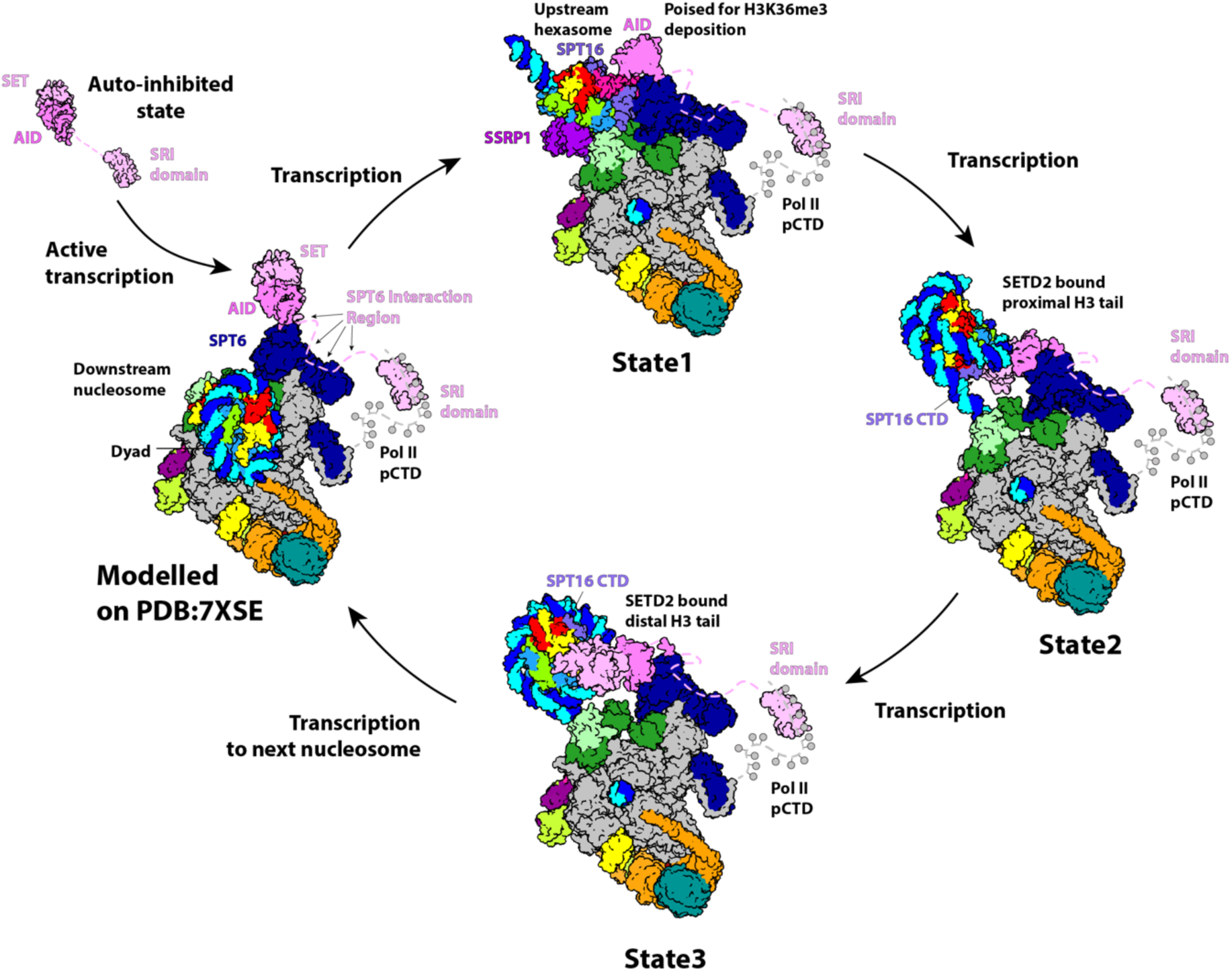
Model for co-transcriptional SPT6 mediated H3K36me3 deposition. SETD2 auto-inhibition is release by bind both the phosphorylated CTD and SPT6 DLD of elongating Pol II. Pol II unwraps the downstream nucleosome during passage with SETD2 bound STP6 waiting nucleosome transfer. FACT mediates nucleosome transfer upstream. Binding of the catalytic SET domain to the transferred hexasome is occluded by FACT and is in a poised state. Deposition of final H2A–H2B dimer allows FACT dissociation from Dyad and SET domain to methylate proximal H3 tail. Unwrapped nucleosome state stabilized by SPT16 CTD. Further transcription rewraps the proximal histones. SETD2 repositions to distal H3 tail whilst remaining in proximity to SPT6 DLD.

In summary, our results provide the first molecular mechanism for co-transcriptional histone modification. Our mechanism resembles that of co-transcriptional RNA transcript modification during pre-mRNA capping and splicing, which also involve interactions with Pol II-associated general elongation factors^70–72^. We note that histone tail modifications other than H3K36me3 are also introduced co-transcriptionally^73^, including ubiquitylation of histone H2B at residue lysine-120 (H2BK120ub1), tri-methylation of histone H3 at residue lysine-4 (H3K4me3) and methylation of H3 at residue lysine-39 (H3K39me). Some of the enzymes required for these modifications interact with the phosphorylated CTD or with Pol II elongation factors^17,19,74–76^. Whether these histone modifications are also introduced after nucleosome transfer to the wake of transcribing Pol II, and which interactions underlie their regulation remains to be investigated in the future.

## Author Contributions

J.L.W., M.O., and P.C. conceived and planned the study; J.L.W., and M.O. designed and performed *in vitro* experiments with U.N. J.L.W., and M.O performed cryo-EM experiments and interpreted data with input from C.D. J.L.W., M.O., U.N., and C.O. purified proteins and nucleic acids. A.Z. conducted preliminary experiments. O.D analyzed MS experiments under H.U. supervision. P.C. acquired funding and supervised the study. J.L.W. and M.O. wrote the original manuscript draft; P.C. reviewed and edited the manuscript, with input from all authors.

## Acknowledgments

We thank U. Neef, P. Rus and T. Schulz for maintaining insect cell facility and pig thymus stocks, C. Dienemann and U. Steuerwald for maintaining the cryo-EM facility. O. Geintzer for maintaining the radioactivity laboratory and discussions of the data. M. Lidschreiber and K. Žumer for providing critical feedback on the manuscript and members of the P.C. laboratory for general discussions of the data.

## Extended Data Figures

**Extended Data Fig. 1:**
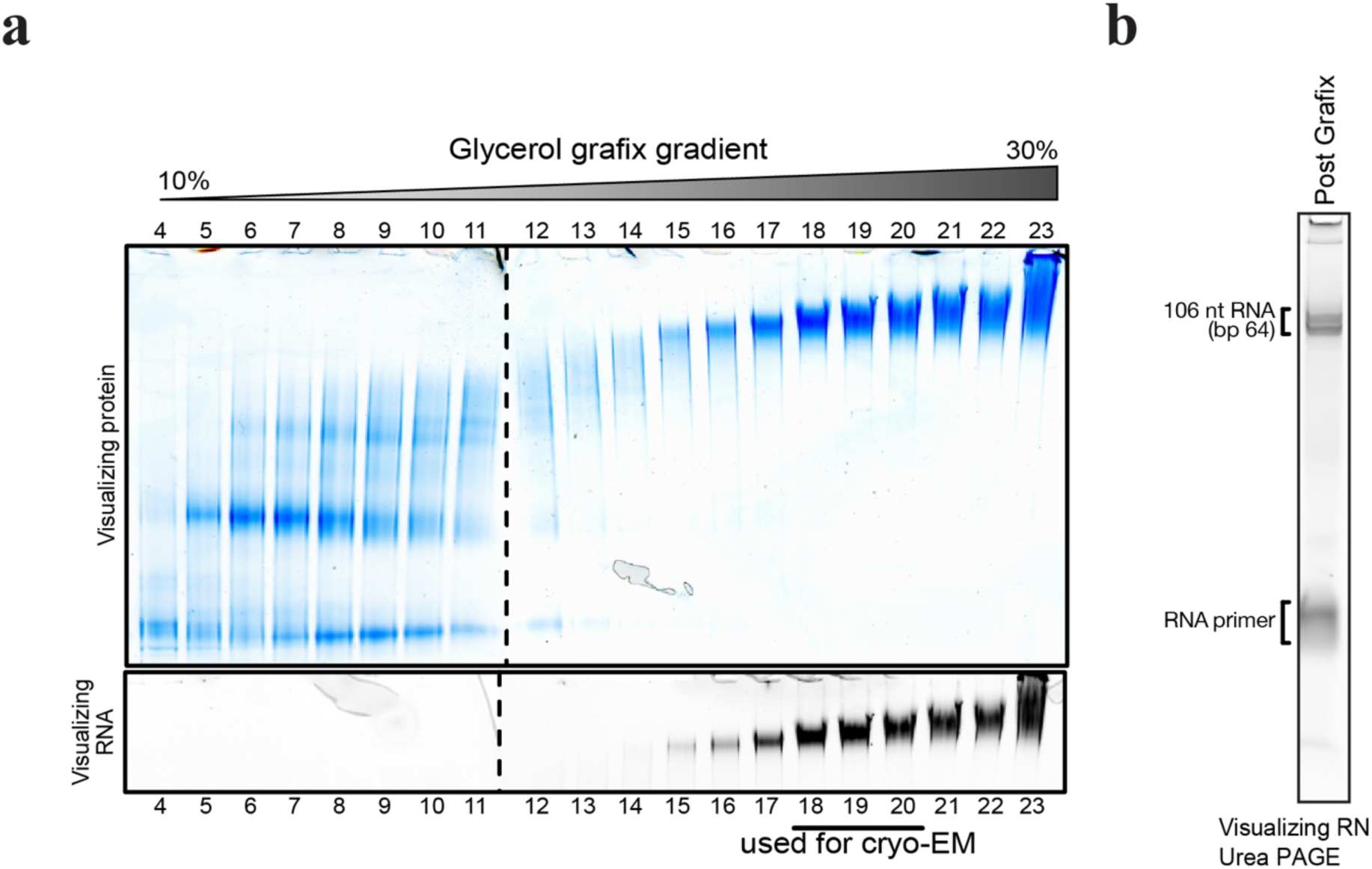
Cryo-EM sample preparation of State 1 complex. **a**, Native PAGE gel of the 10-30% Grafix glycerol gradient fractions of the methylation competent elongation complex transcribed to nucleosomal bp 64. In the top gel image, the peptides are visualized using Coomassie staining. The bottom gel image visualizes the 5’Cy5 labelled RNA. Fractions used for cryo-EM analysis are indicated. **b**, Denaturing PAGE of complex formation for cryo-EM analysis to verify RNA extension to nucleosomal bp 64. The 5’Cy5 label of the RNA was visualized.

**Extended Data Fig. 2:**
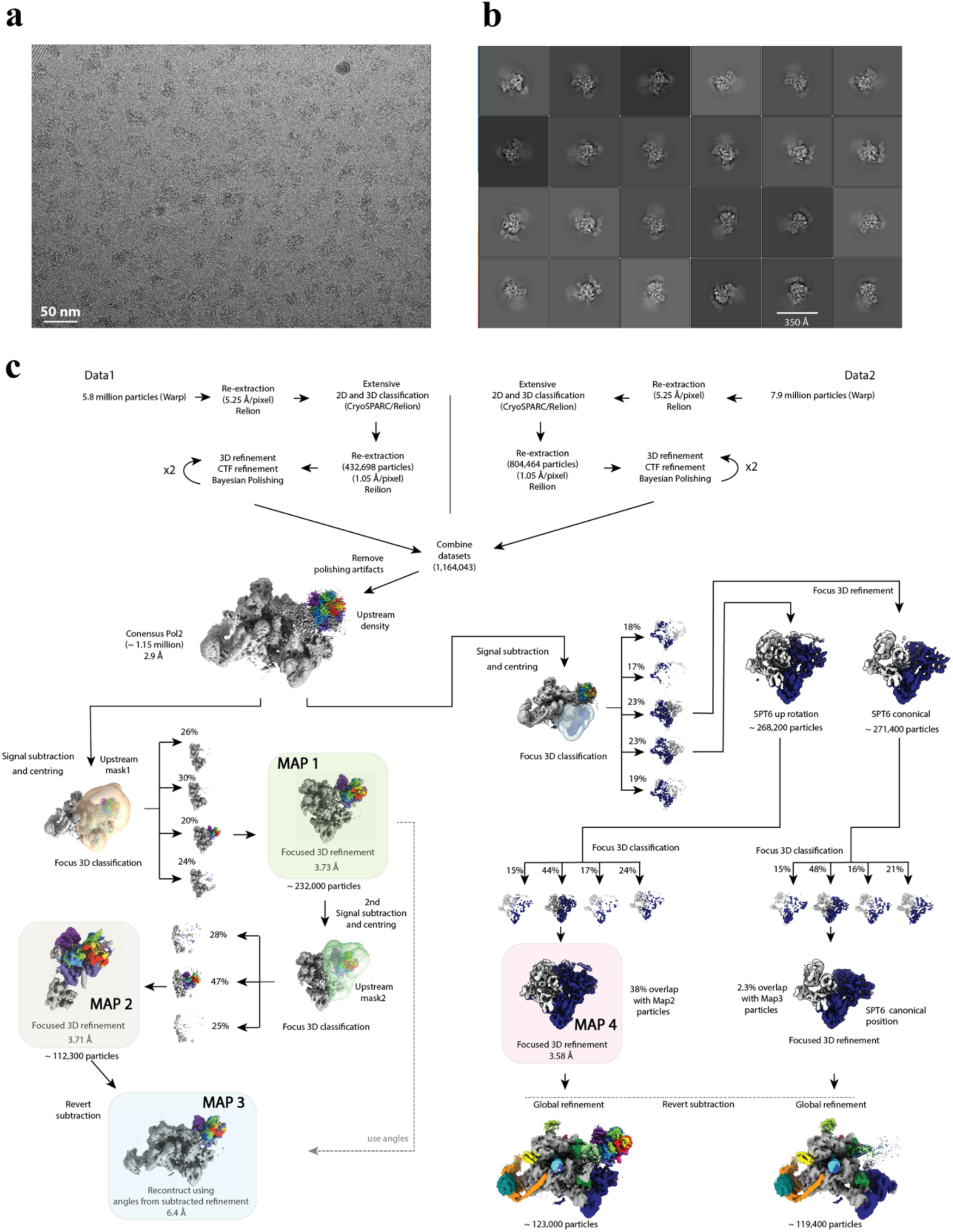
Data acquisition and processing of State 1 complex. **a**, Representative micrograph of data collection with scale bar. **b**, Representative 2D classes of mammalian activated Elongation complex obtained during junk removal. **c**, Processing and classification strategy employed to classify the methylation competent elongation complex from obtained micrographs. Major steps are described within the figure. Coulomb maps used to build models are highlighted.

**Extended Data Fig. 3:**
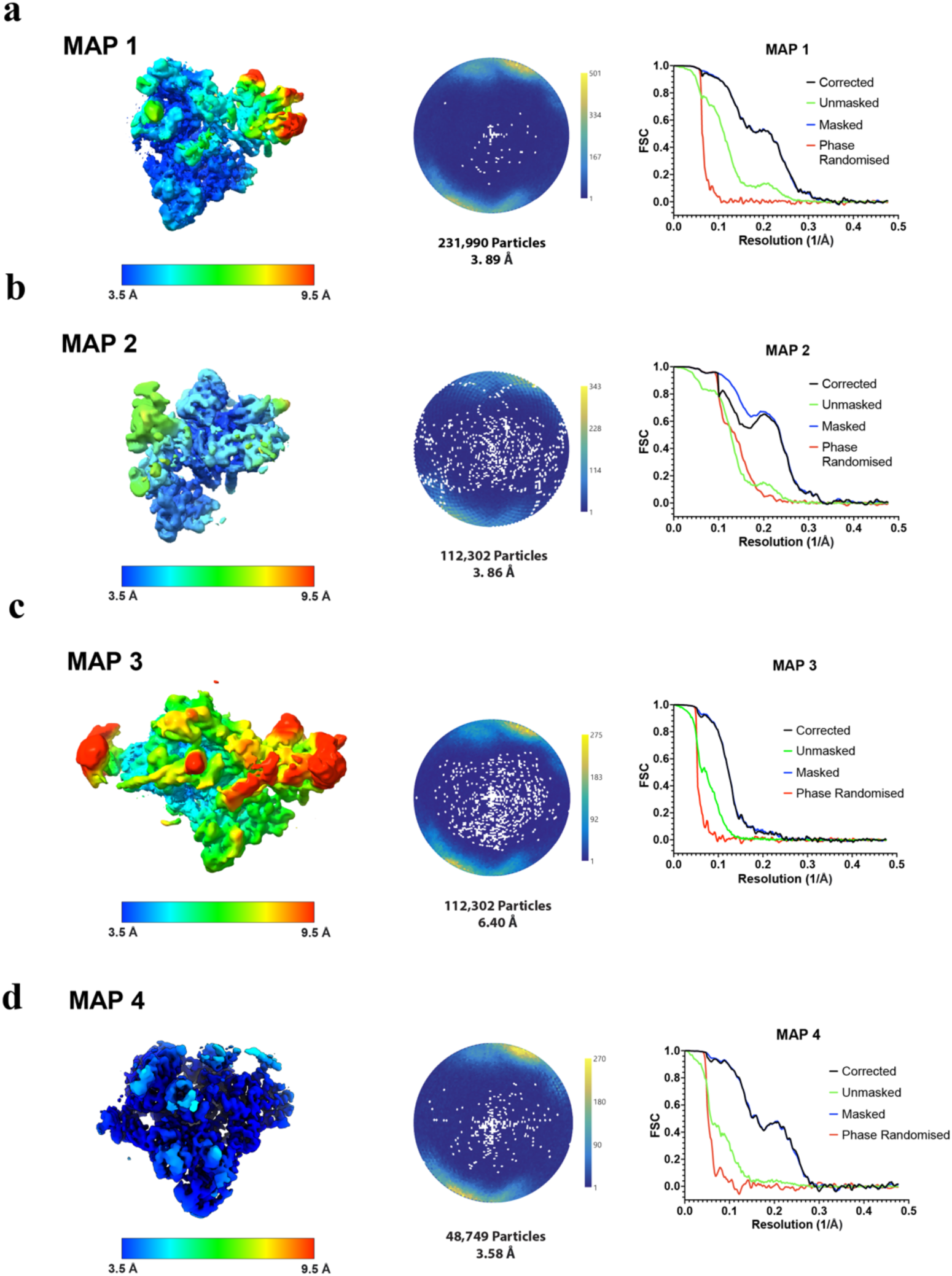
Cryo-EM data metrics of State 1 complex. Reconstructions obtained from State 1 dataset coloured by their local resolution as estimated using RELION. Total particle count, the global resolution estimate using the Fourier shell correlation (FSC) = 0.143 criterion are given together with an angular distribution plot. MAP 1 **a**, MAP 2 **b**, MAP 3 **c**, Map 4 **d**,.

**Extended Data Fig. 4:**
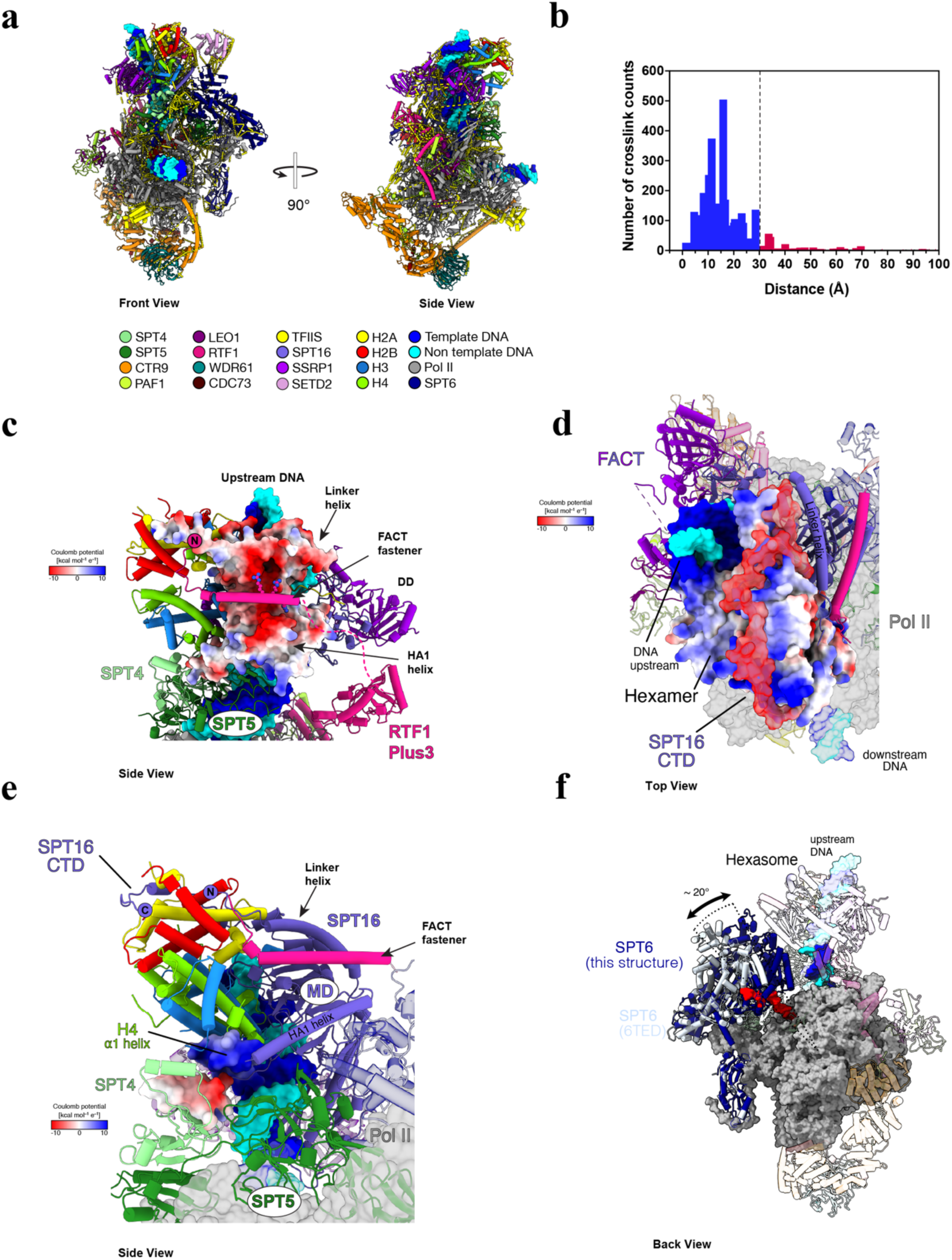
Chemical Xlinking mass spec and State 1 interactions. **a**, Front and Side view of State 1 with mapped BS3 crosslinks (cutoff >30 Å). **b**, Histogram of crosslink counts and their distances between Nζ pairs that were mapped onto the model. In case the side chain was not modelled, C⍺ pairs were used. 9 crosslinks with a distance longer than 100 Å were excluded from the histogram. **c**, SPT16 CTD binds exposed histones. SPT16 is shown by its Coulombic surface potential. Arg 289, Arg 292 and Arg 295 in RTF1 FACT “fastener” shown as spheres. **d**, Top view of upstream hexasome in State 1. Histones and SPT16 CTD are shown by their Coulombic surface potential. **e**, Interaction of DSIF with histones and SPT16. SPT4 interacts with ⍺1 helix of H4 (both partially presented by their Coulombic surface potential). SPT5 contacts SPT16’s HA1 helix and MD. **f**, SPT6 rearranges to contact the upstream hexasome bound by FACT. Structural comparison of the activated elongation complex (PDB: 6TED – coloured light steel blue) ^10^ reveals significant rotation (∼20°) and rearrangement of SPT6 to contact upstream hexasome. State 1 SPT6 shown in dark blue, SPT6. Other elongation factors and the upstream hexasome with FACT are shown transparent.

**Extended Data Fig. 5:**
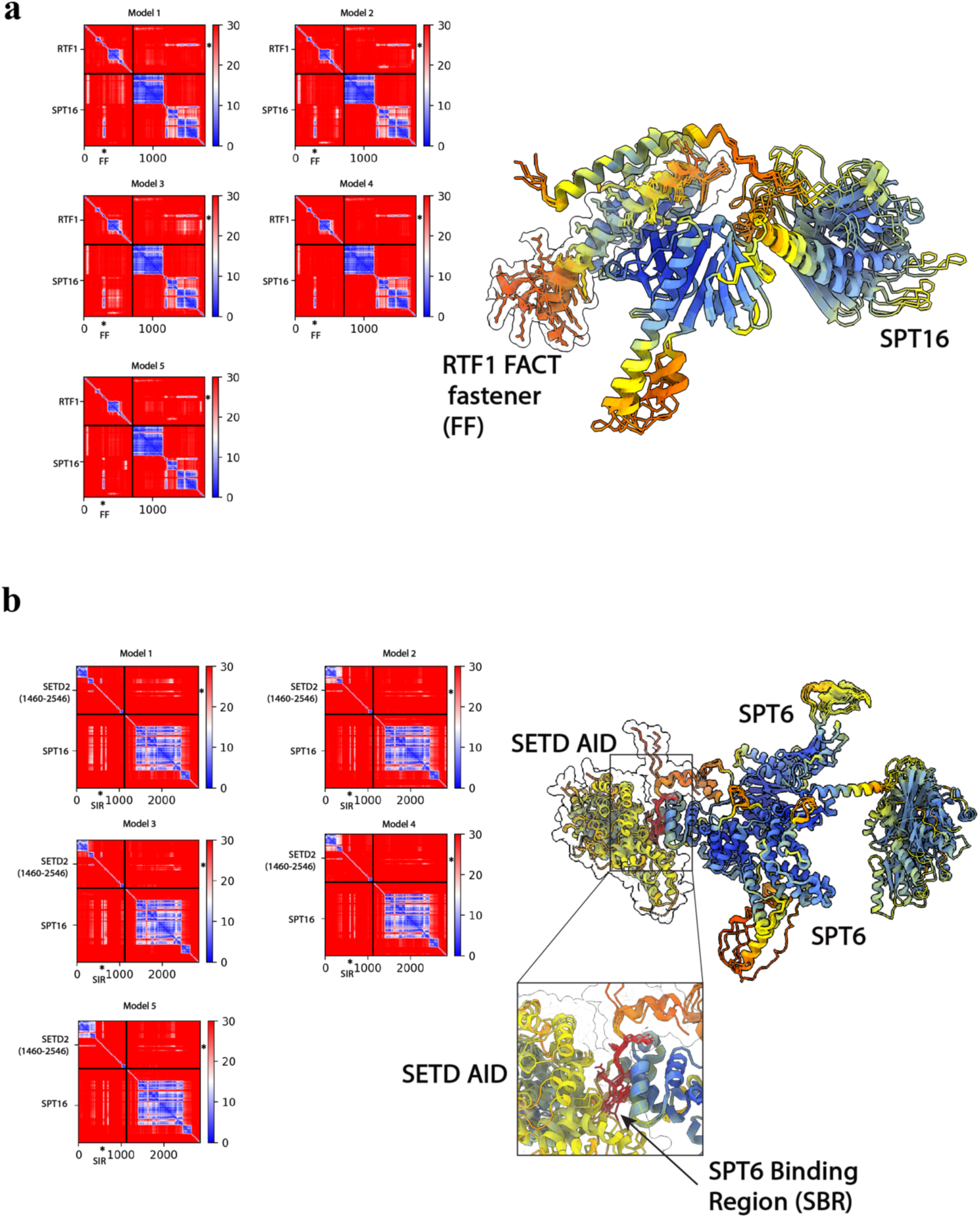
ColabFold prediction of protein interactions. **a**, Left – Predicted Aligned Error (PAE) of full-length RTF1 and SPT16. Predicted FACT “fastener” interaction marked as an asterisk. Right – Overlay of 5 predicted models of SPT16 (residues 466-929 shown) and RTF1 (residues 266-304 shown). Cartoon colored by pLDDT confidence. **b**, Left – Predicted Aligned Error (PAE) of full-length SETD2 (residues 1460-2546) and SPT6. Predicted SBR interaction marked as an asterisk. Right – Overlay of 5 predicted models of SPT6 (residues 1731-1852,2002-2059 shown) and SPT6 (residues 266-1521 shows). Cartoon colored by pLDDT confidence.

**Extended Data Fig. 6:**
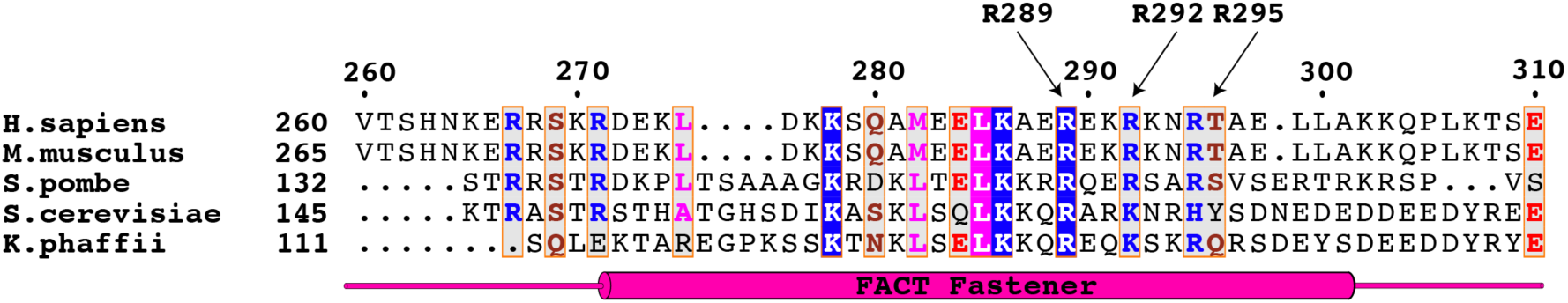
RTF1 FACT “fastener” sequence alignment. Sequence alignment of the FACT “fastener” from RTF1 generated using ClustalOmega^77^ and visualized in ESPript^78^. *Homo sapiens*, *Mus musculus*, *Schizosaccharomyces pombe*, *Saccharomyces cerevisiae* and *Komagataella phaffii* were aligned and compared. The FACT “fastener” of RTF1 and its positive charge is highly conserved. Arg289, Arg292 and Arg295 of the FACT “fastener” are highlighted by arrows.

**Extended Data Fig. 7:**
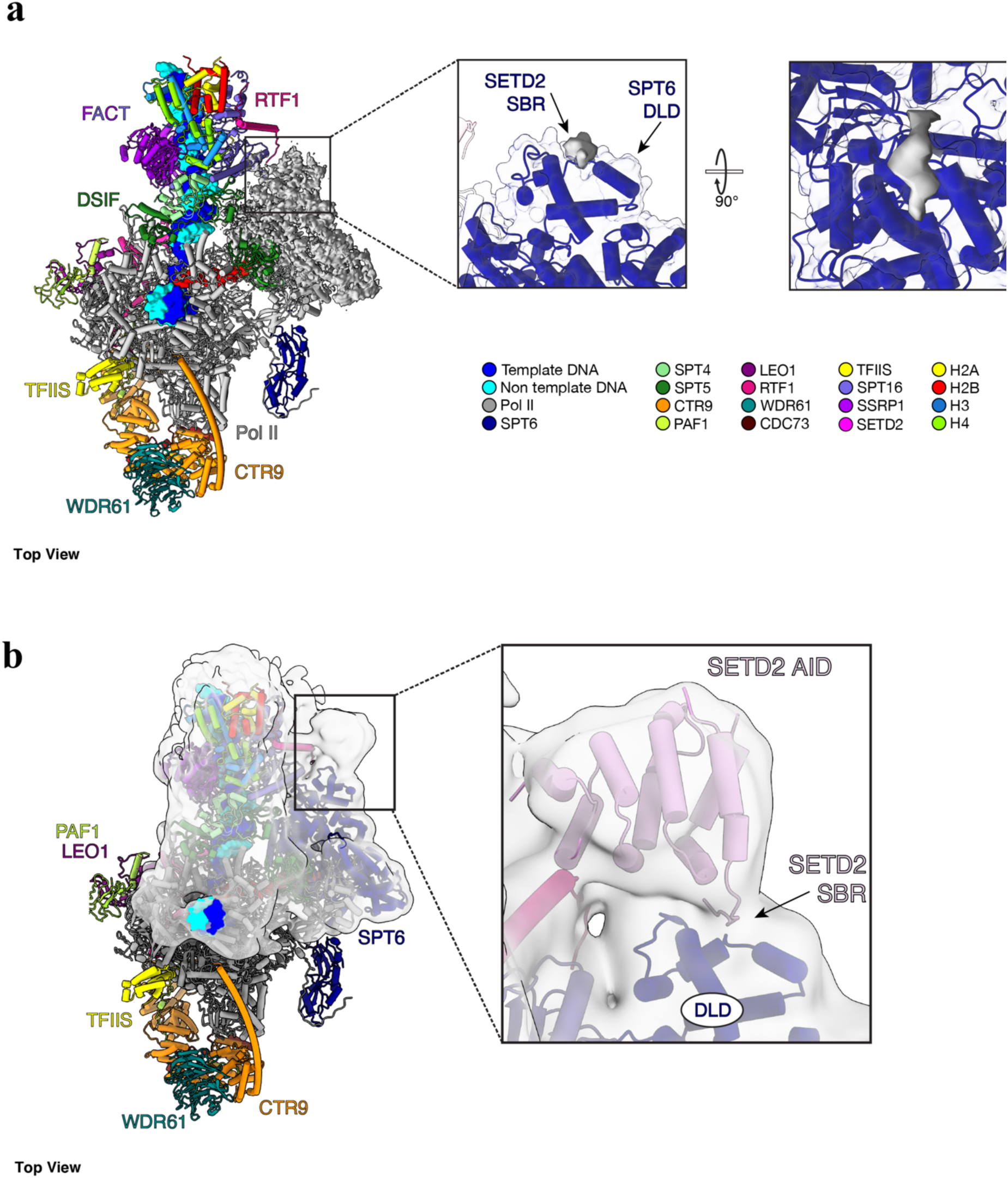
SETD2 binds SPT6 DLD. **a**, Top view of State 1 with cryo-EM density of MAP 3 (Extended Data Fig. 3b) displayed in grey. Cryo-EM density for the modelled region the SETD2 SPT6 Interaction Region shown in zoomed panels. **b**, Top view of State 1 with the Gaussian filtered (σ = 2) cryo-EM density of MAP 1 (Extended Data Fig. 3b) shown in grey. Alphafold2 model for SETD2 AID shown in panel and rigid body fitted into density. Interaction between SPT6 DLD and SETD2 modelled based of ColabFold predictions (Extended Data Fig. 5)

**Extended Data Fig. 8:**
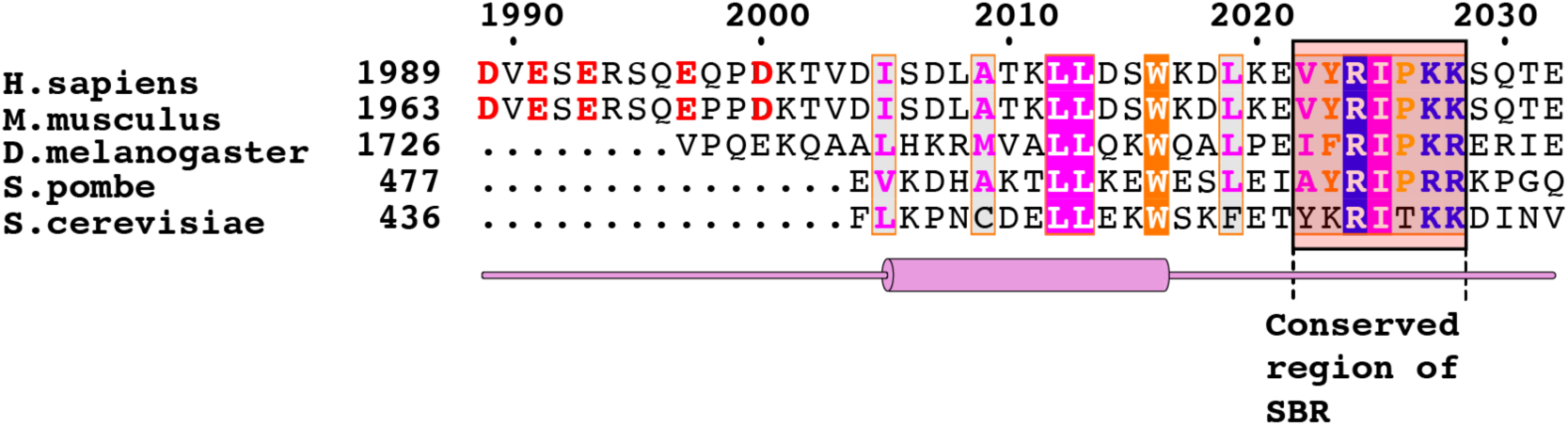
SETD2 SPT6 Interaction Region sequence alignment. Sequence alignment of the SETD2 SPT6 Interaction Region generated using ClustalOmega ^77^ and visualized in ESPript ^78^. *Homo sapiens*, *Mus musculus*, *Schizosaccharomyces pombe*, *Saccharomyces cerevisiae* and *Drosophila melanogaster* were aligned and compared. Resides 2022 – 2028 are highly conserved.

**Extended Data Fig. 9:**
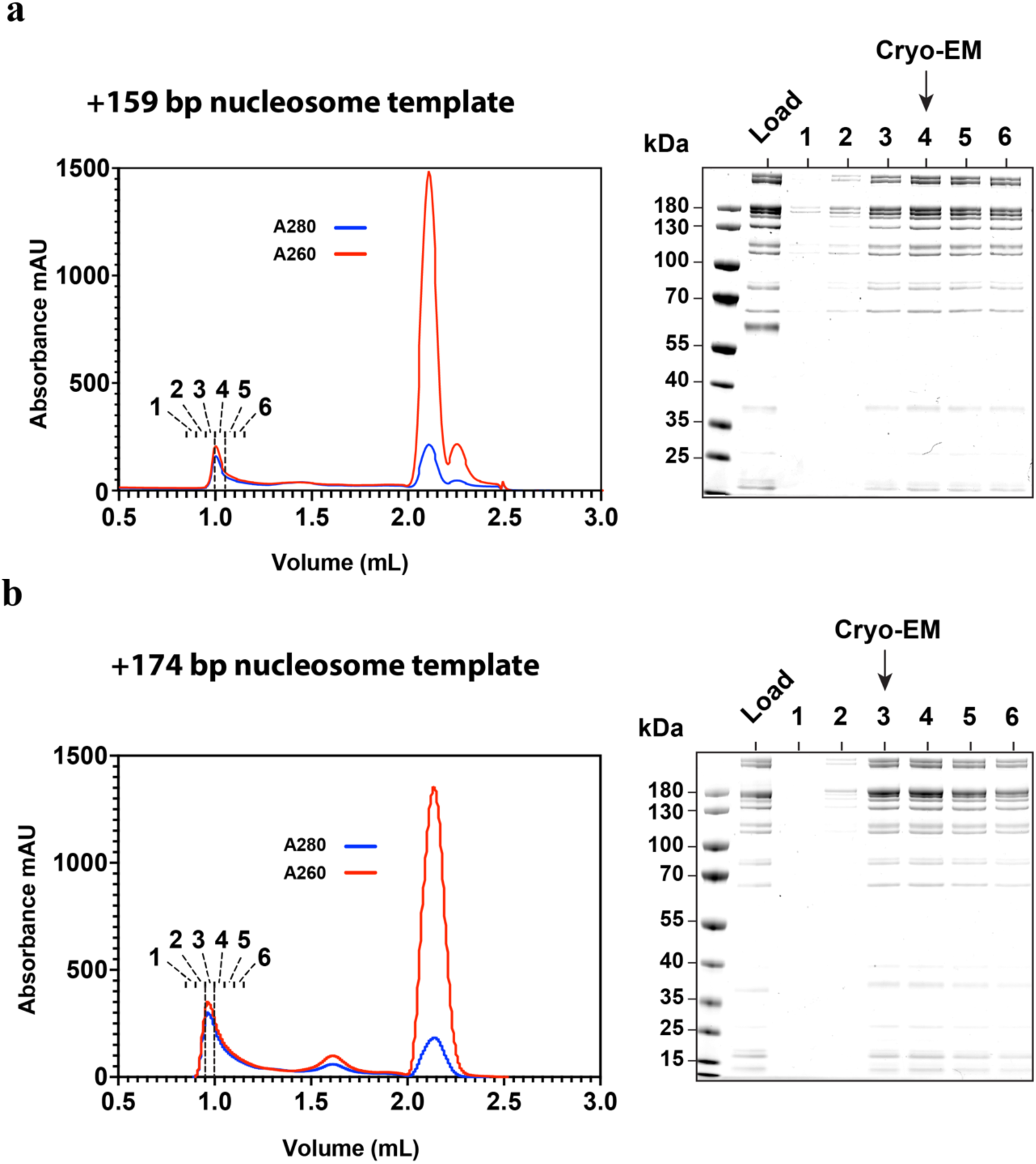
Cryo-EM sample purification of State 2 and State 3 complexes. **a**, Left – Chromatogram of Superose 6 Increase (3.2/300) purification of methylation competent elongation complex bound to a ligated nucleosome template that would represent transcription to bp 159 into the nucleosome template. Right – SDS-PAGE analysis of peak fractions. Indicated fraction used for cryo-EM analysis. **b**, Left – Chromatogram of Superose 6 Increase (3.2/300) purification of methylation competent elongation complex bound to a ligated nucleosome template that would represent transcription to bp 174 into the nucleosome template. Right-SDS-PAGE analysis of peak fractions. Indicated fraction used for cryo-EM analysis.

**Extended Data Fig. 10:**
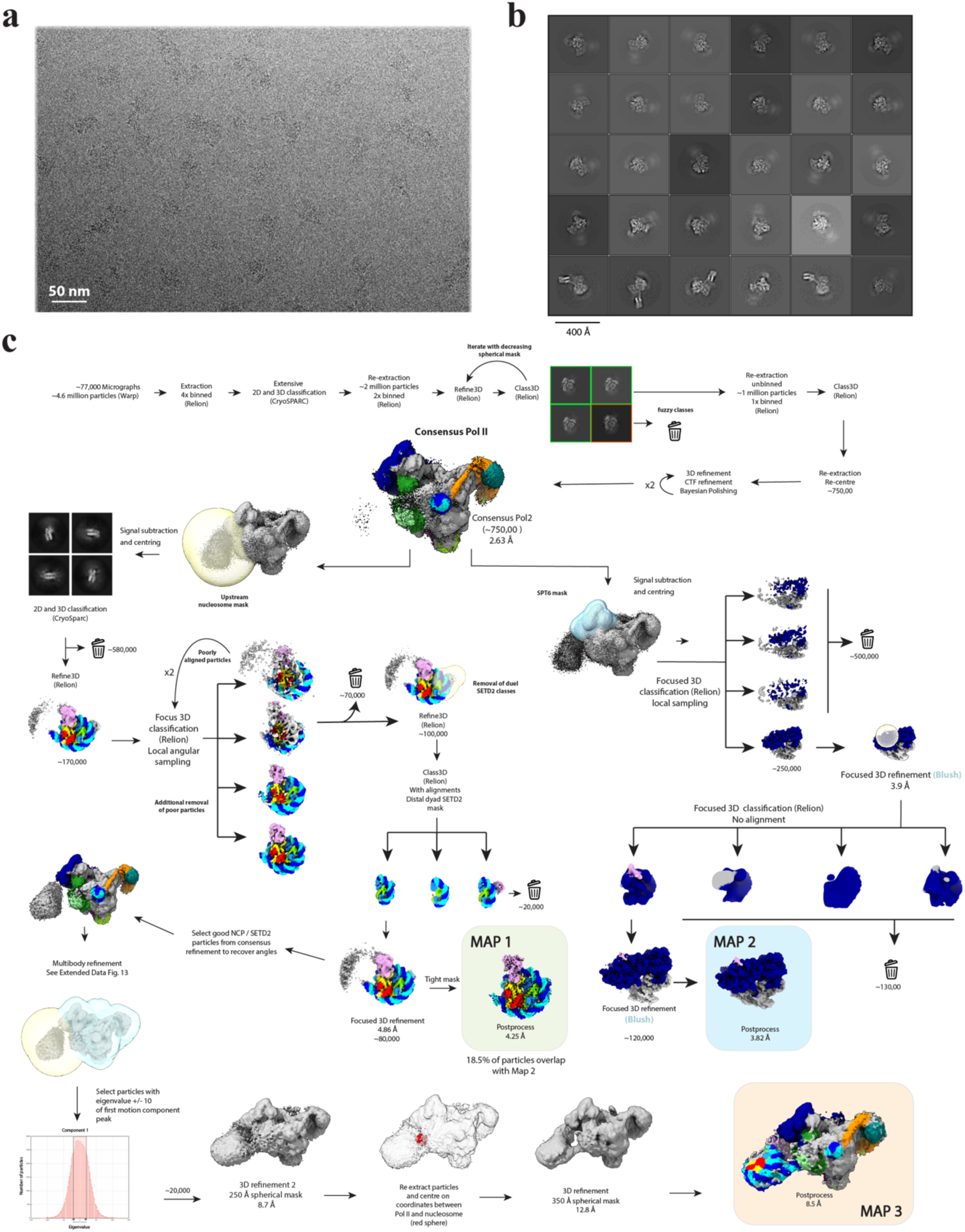
Data acquisition and processing of State 2 complex. **a**, Representative micrograph of data collection with scale bar. **b**, Representative 2D classes of mammalian activated elongation complex obtained during junk removal. **c**, Processing and classification strategy employed to classify the methylation competent elongation complex from obtained micrographs. Major steps are described within the figure. Coulomb maps used to build models are highlighted.

**Extended Data Fig. 11:**
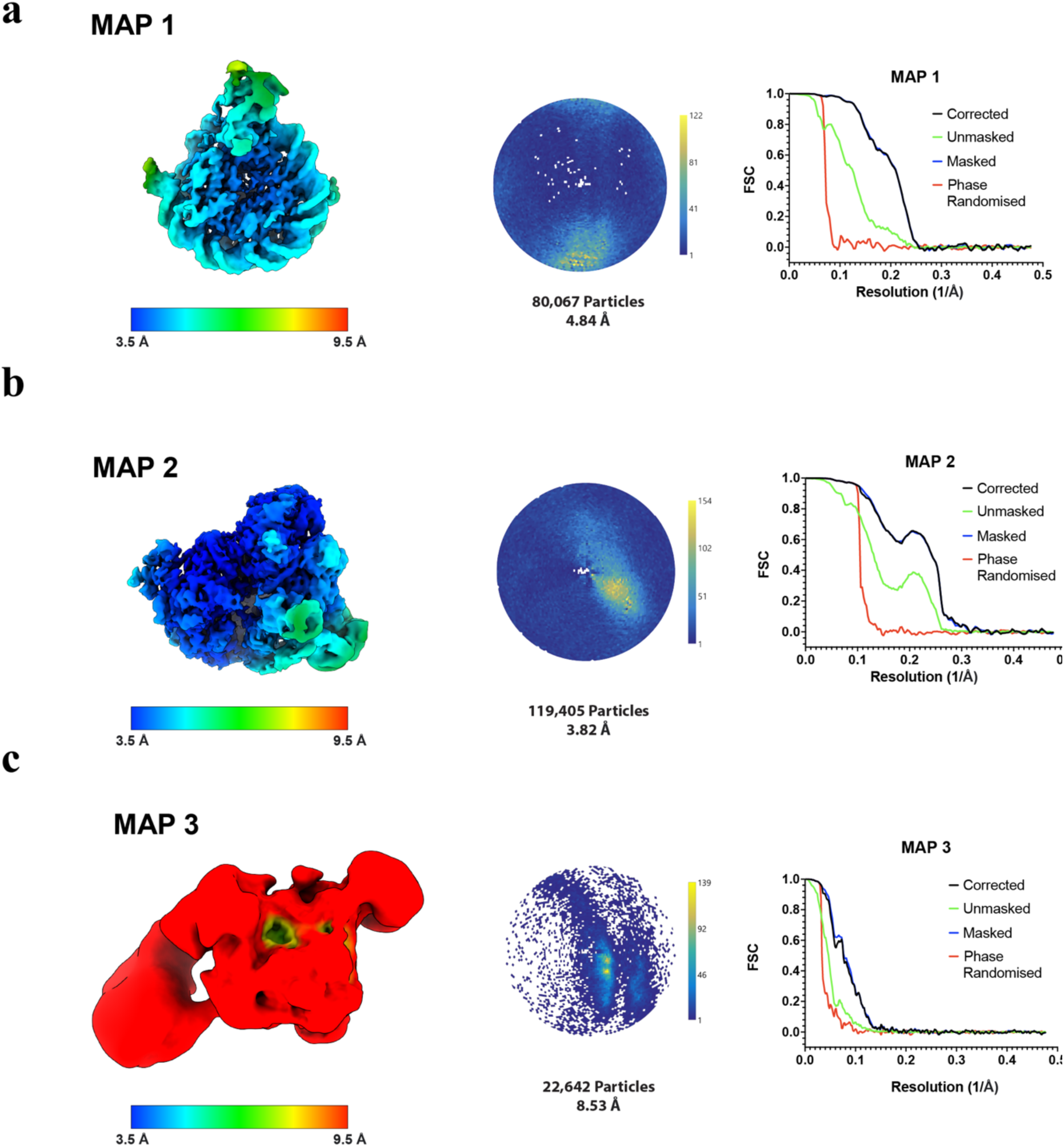
Cryo-EM data metrics of State 2 complex. Reconstructions obtained from State 1 dataset coloured by their local resolution as estimated using RELION. Total particle count, the global resolution estimate using the Fourier shell correlation (FSC) = 0.143 criterion are given together with an angular distribution plot **a**, MAP 1 **b**, MAP 2 **c**, MAP 3.

**Extended Data Fig. 12:**
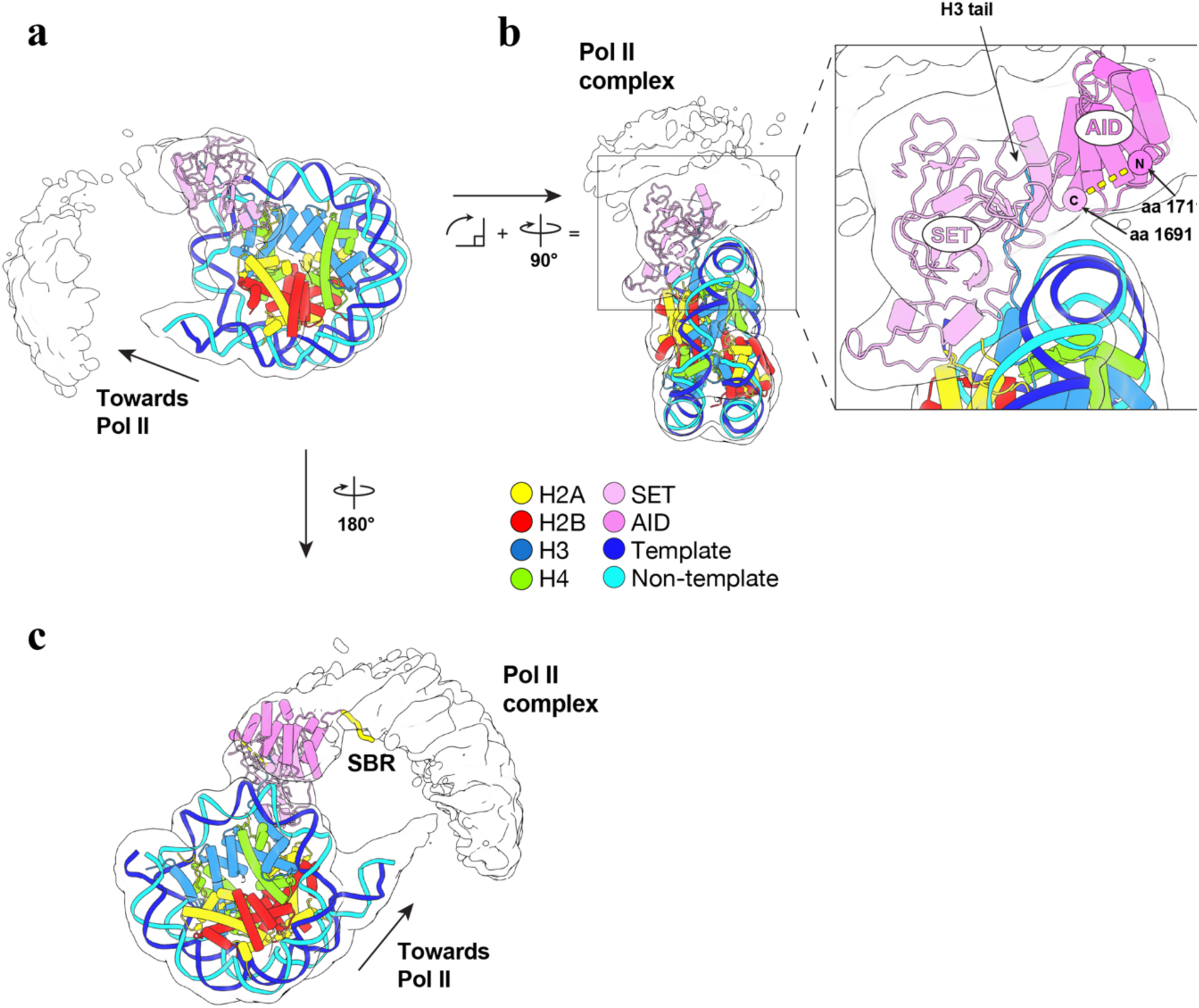
SETD2 bound nucleosome in State 2 complex. **a**, View of State 2 SETD2 bound nucleosome with Gaussian filtered (σ = 2) cryo-EM density of MAP 1 (Extended Data Fig. 11a) displayed in grey. **b**, Alphafold2 model for SETD2 AID manually fitted into density and arranged to minimize distance between N and C terminal. **c**, Alternate view of (a, b) with conserved residues of SBR shown as yellow tube.

**Extended Data Fig. 13:**
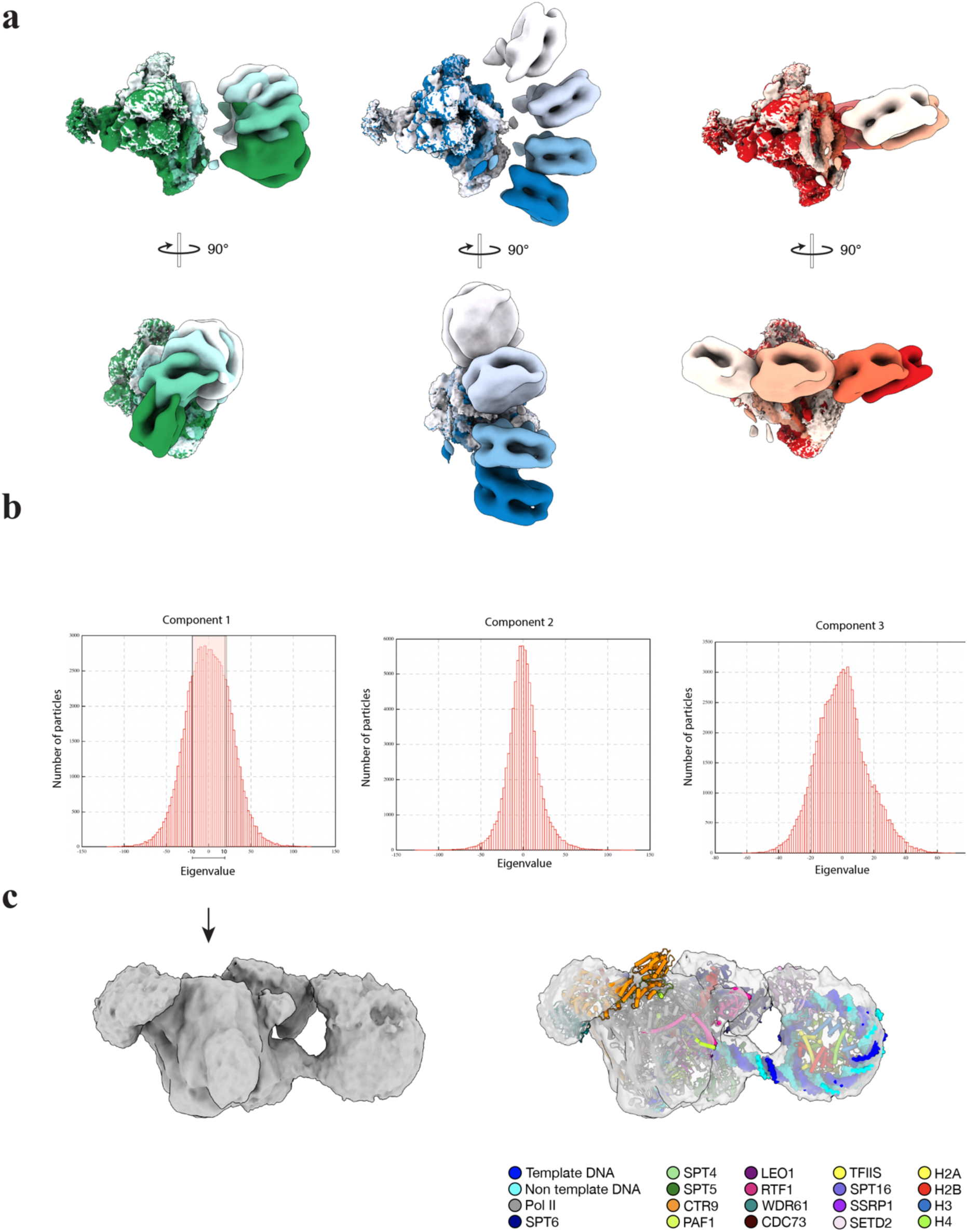
Multibody refinement analysis of State 2. **a**, Selected maps showing the extent of the first 3 principal motion components from multibody refinement analysis of State 2 are shown in green, blue and red respectively. **b**, Eigenvalues for the first 3 principal motion component. Selected values of component 1 highlighted. c, Refinement of selected particles from component 1 (after re-centering) with fitted model of State 2 complex.

**Extended Data Fig. 14:**
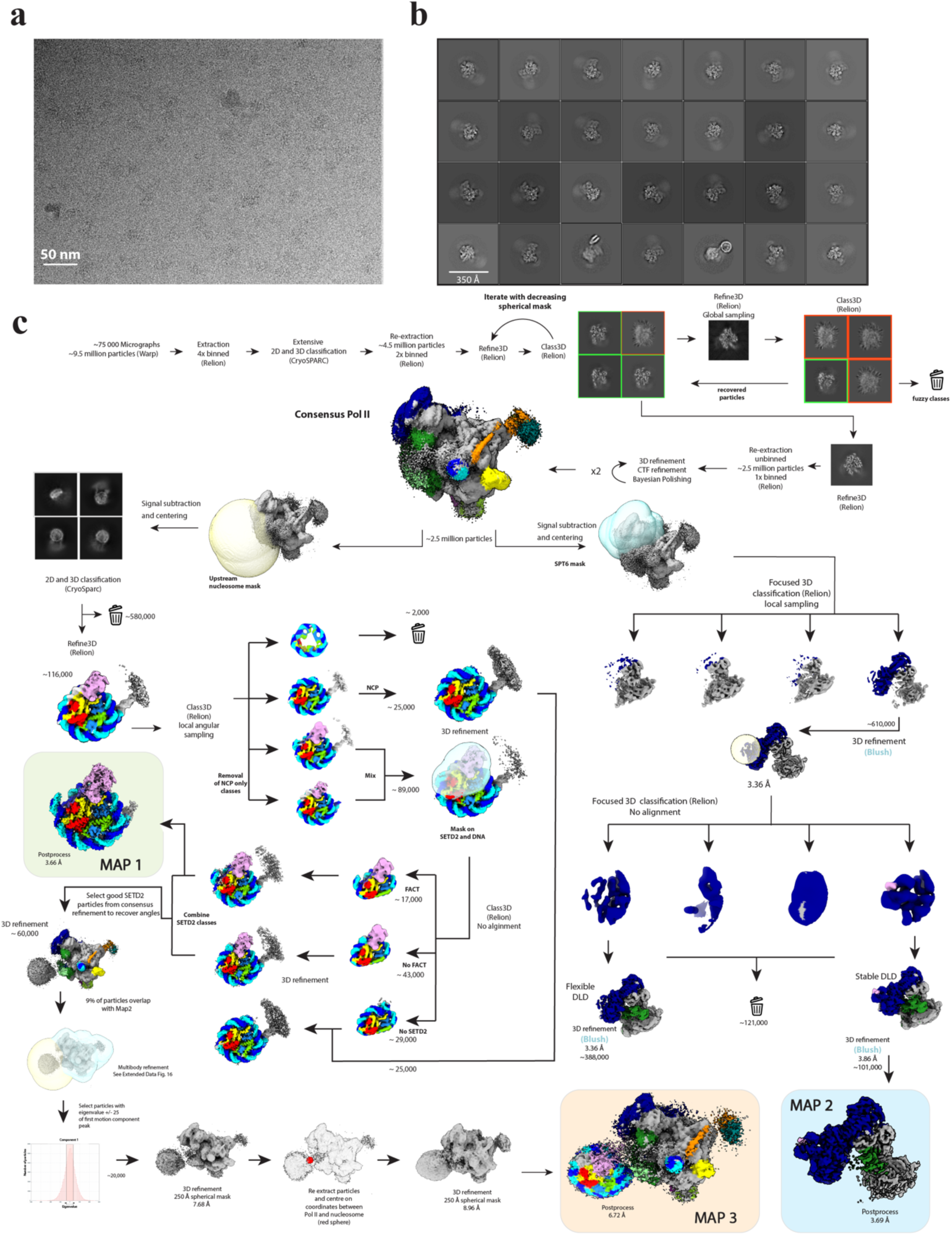
Data acquisition and processing of State 3 complex. **a**, Representative micrograph of data collection with scale bar **b**, Representative 2D classes of mammalian activated elongation complex obtained during junk removal. **c**, Processing and classification strategy employed to classify the methylation competent elongation complex from obtained micrographs. Major steps are described within the figure. Coulomb maps used to build models are highlighted.

**Extended Data Fig. 15:**
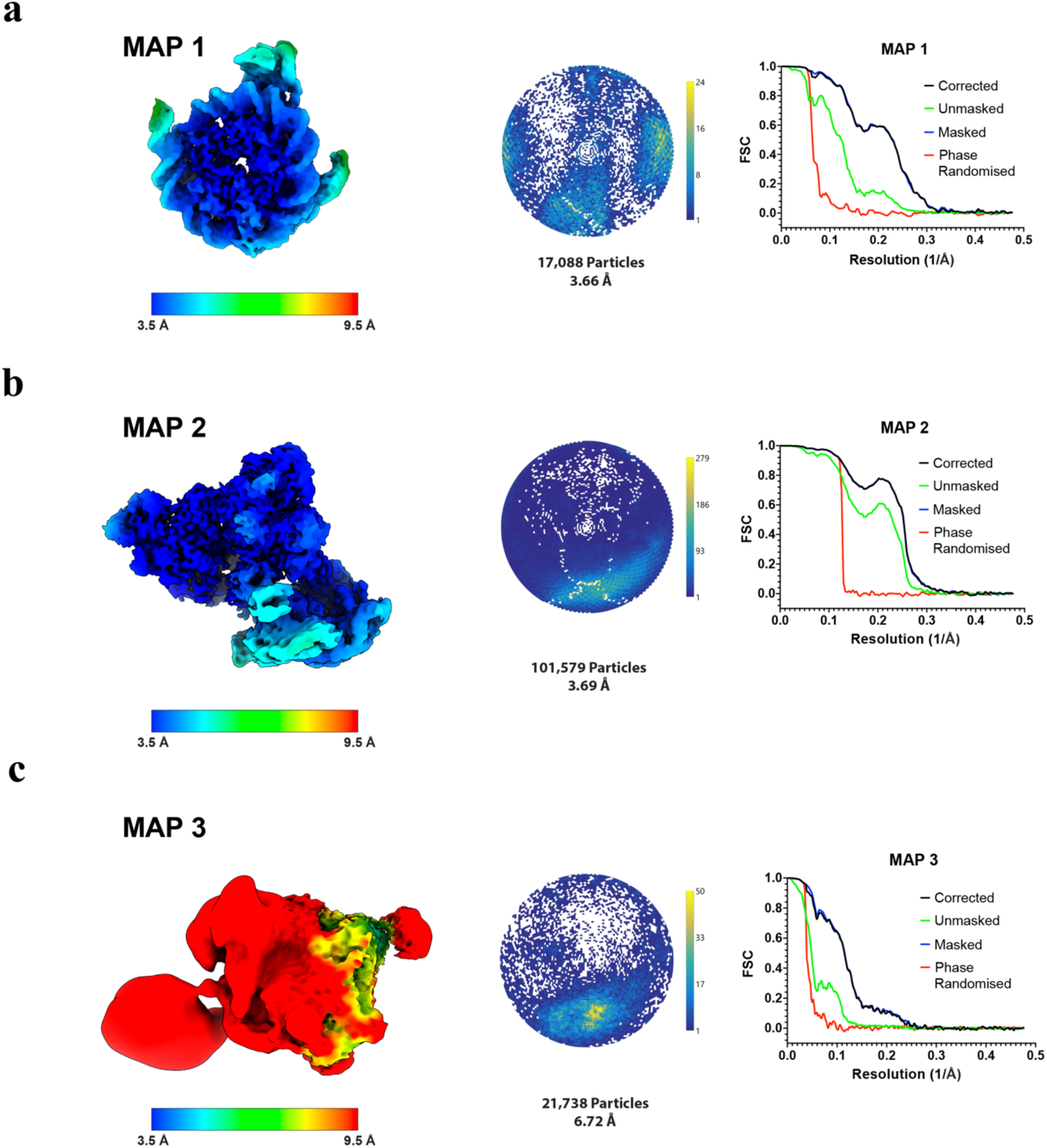
Cryo-EM data metrics of State 3 complex. Reconstructions obtained from State 1 dataset coloured by their local resolution as estimated using RELION. Total particle count, the global resolution estimate using the Fourier shell correlation (FSC) = 0.143 criterion are given together with an angular distribution plot. **a**, MAP 1 **b**, MAP 2 **c**, MAP 3.

**Extended Data Fig. 16:**
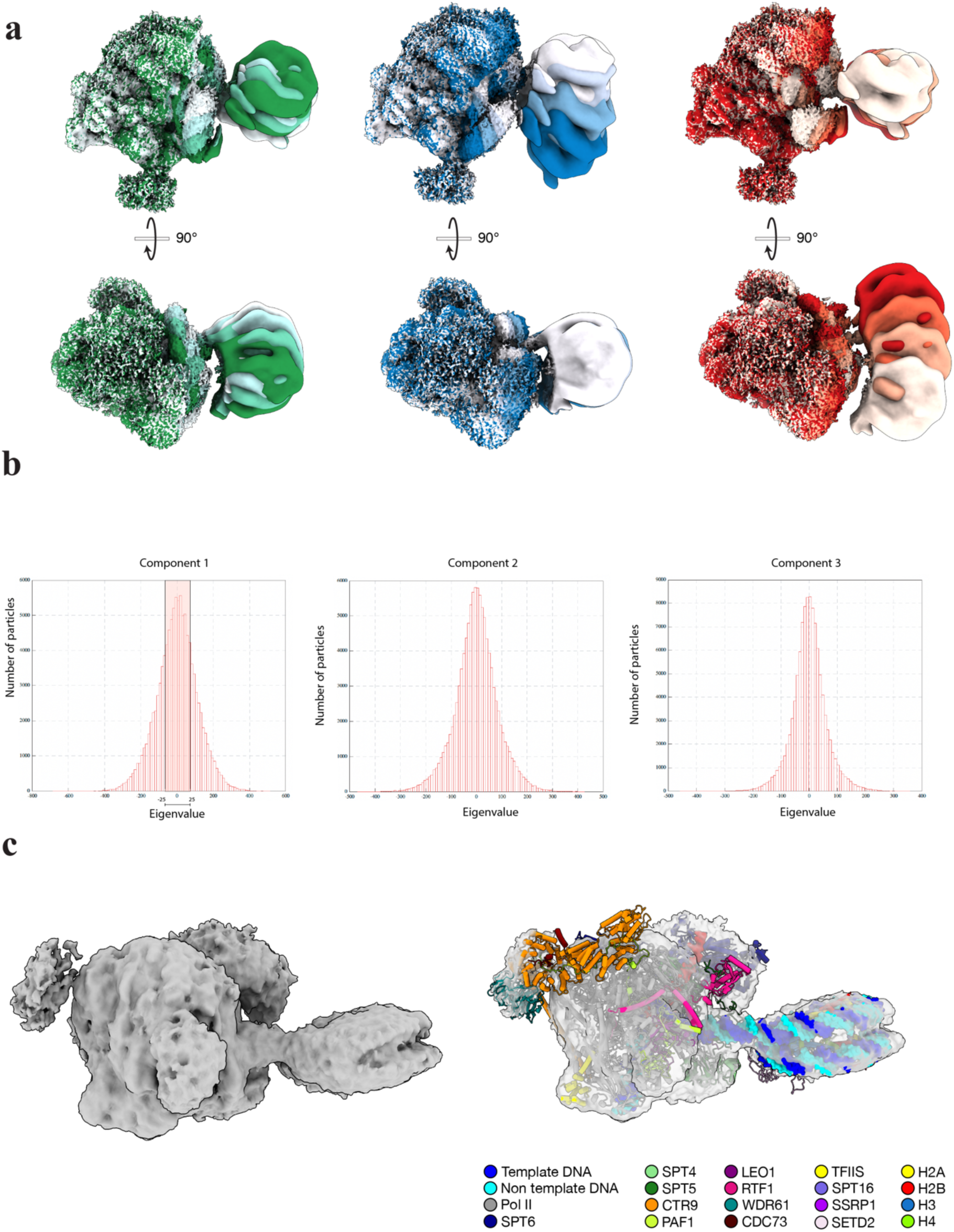
Multibody refinement analysis of State 3. **a**, Selected maps showing the extent of the first 3 principal motion components from multibody refinement analysis of State 3 are shown in green, blue and red respectively. **b**, Eigenvalues for the first 3 principal motion component. Selected values of component 1 highlighted. c, Refinement of selected particles (after re-centering) from component 1 with fitted model of State 2 complex.

## Methods

### Resource Availability

Requests for materials should be directed and will be fulfilled by the corresponding author, Patrick Cramer.

### Materials Availability

Materials are available from the corresponding author Patrick Cramer upon request under a material transfer agreement with the Max Planck Society.

### Method Details

#### Preparation of protein components

*Sus scrofa* RNA polymerase II was extracted from 0.5 – 2 kg pig thymus. *Homo sapiens* SPT6, PAF1c, RTF1, IWS1, SETD2, AKT3, and FACT were expressed in *Trichoplusia ni* Hi5 cells and harvested after approx. 72 hours. H. sapiens TFIIS, DSIF, and histones H2A, H2B, H3, H3K36M, and H4 were expressed for 3 h in BL21 (DE3) RIL *Escherichia coli* cells. *S. scrofa* Pol II and *H. sapiens* DSIF, TFIIS^7^, SPT6, PAF1c, RTF1^10,12^, FACT^33^, human histones H2A, H2B, H3, H3K36M, and H4^79,80^ were purified as previously described. *H. sapiens* IWS1 was purified from 750 mL of Hi5 expression. All steps were performed at 4°C. The cell pellet was resuspended and lysed by sonication in lysis buffer (300 mM NaCl, 20 mM Na·HEPES pH 7.4 (@ 20 °C), 30 mM imidazole, 10% (v/v) glycerol, 1 mM DTT, 0.284 µg mL^−1^ leupeptin, 1.37 µg mL^−1^ pepstatin A, 0.17 mg mL^−1^ PMSF, and 0.33 mg mL^−1^ benzamidine). Cell depris were cleared by centrifugation and the supernatant filtered through 0.8 µM syringe filter. The sample was applied to two pre-equilibrated 5 mL HisTrap HP column (Cytivia). The column was washed with 10 CV lysis buffer, 5 CV high salt buffer (1000 mM NaCl, 20 mM Na·HEPES pH 7.4 (@ 20 °C), 30 mM imidazole, 10% (v/v) glycerol, 1 mM DTT, 0.284 µg mL^−1^ leupeptin, 1.37 µg mL^−1^ pepstatin A, 0.17 mg mL^−1^ PMSF, and 0.33 mg mL^−1^ benzamidine), and 5 CV low salt buffer (150 mM NaCl, 20 mM Na·HEPES pH 7.4 (@ 20 °C), 30 mM imidazole, 10% (v/v) glycerol, 1 mM DTT, 0.284 µg mL^−1^ leupeptin, 1.37 µg mL^−1^ pepstatin A, 0.17 mg mL^−1^ PMSF, and 0.33 mg mL^−1^ benzamidine). A low salt equilibrated 5 mL HiTrap Q HP column (Cytivia) was attached to the base of the HisTrap column. The HisTrap column was developed with a 3 CV long gradient in nickel elution buffer (150 mM NaCl, 20 mM Na·HEPES pH 7.4 (@ 20 °C), 500 mM imidazole, 10% (v/v) glycerol, 1 mM DTT, 0.284 µg mL^−1^ leupeptin, 1.37 µg mL^−1^ pepstatin A, 0.17 mg mL^−1^ PMSF and 0.33 mg mL^−1^ benzamidine), before the column was removed. The remaining HiTrap Q column was washed with 5 CV low salt buffer before it was developed with a 9 CV high salt buffer gradient. Peak fractions were analyzed by SDS-PAGE and Coomassie staining. Fractions containing IWS1 were pooled and mixed with TEV protease, lambda protein phosphatase, and 1 mM MnCl2. The sample was dialysed overnight using Snake Skin dialysis tubing (7 kDa molecular weight cutoff) (Thermo Scientific™) against 1 L lysis buffer containing 1 mM MnCl2. The sample was then applied to two pre-equlibrated 5 mL HisTrap HP column. The flow-through was collected and concentrated in a pre-equilibrated 50kMWCO Amicon Ultra Centrifugal Filters (Milipore). The concentrated sample was then applied to a HiLoad 16/600 Superdex 200pg column equilibrated in SEC buffer (300 mM NaCl, 20 mM Na·HEPES pH 7.4 (@ 20 °C), 10% (v/v) glycerol, 1 mM DTT). Peak fractions were analyzed by SDS-PAGE and Coomassie staining. Fractions containing pure IWS1 were pooled, concentrated in a pre-equilibrated 50kMWCO Amicon Ultra Centrifugal Filters (Milipore), and snap frozen in liquid nitrogen. The protein was stored at –70 °C until use.

SETD2 (residues 1446-2564) expressed in Hi5 insect cells as an 6xHis MBP fusion protein and sonication and clarification were performed as described for IWS1. Clarified lysate was applied to a XK 16/20 column packed with amylose resin. The column was washed with 3 column volumes of lysis buffer (300 mM NaCl, 20 mM Na-HEPES, pH 7.4, 30 mM imidazole, 10% (v/v) glycerol, 1 mM DTT, 0.284 μg ml−1 leupeptin, 1.37 μg ml−l pepstatin A, 0.17 mg ml−1 PMSF and 0.33 mg ml−1 benzamidine) and SETD2 eluted with 2 column volumes of elution buffer (lysis buffer supplemented with 10 mM maltose). Eluted fractions were analyzed by denaturing polyacrylamide gel electrophoresis (SDS-PAGE) and fractions containing SETD2 were pooled and incubated with TEV protease for a minimum of 12 hours. The cleaved sample was applied to two 5 mL HisTrap HP to remove TEV and MBP. The columns were washed with one column volume of lysis buffer and the cleaved SETD2 recovered from the flow-through, concentrated in a 50 kDa MWCO Amicon Ultra Centrifugal Filter (Merck) and applied to HiLoad S200 16/600 pg column equilibrated with gel filtration buffer (300 mM NaCl, 20 mM Na-HEPES, pH 7.4, 10% (v/v) glycerol, 1 mM DTT). Peak fractions were assessed by SDS-PAGE analysis. SETD2 fractions were pooled and concentrated in a 50 kDa MWCO Amicon Ultra Centrifugal Filter (Merck), aliquoted, flash frozen in liquid nitrogen and stored at −70 °C.

Protein truncations and deletions were expressed and purified in identical fashion to WT proteins.

#### Preparation of mononucleosomal and tetranucleosomal constructs

DNA constructs for mononucleosomal templates were generated by 50 mL PCR^6,7,35,80^. The template for amplification were vectors containing the Widom 601 nucleosome positioning sequence with T-less cassette on the template strand to nucleosomal bp 32 or 64. PCR reactions were performed with two primers (forward primer: 5’-GCA GTC CAG TTA CGC TGG AGT C-3’; reverse primer: 5’-ATC AGA ATC CCG GTG CCG-3’). The PCR products were purified by anion exchange, digested by TspRI, and reconstituted with octamer to mononucleosomes. The DNA construct featuring the T-less cassette was used for structural analysis by cryo-EM.

To generate nucleosome templates for cryo-EM studies, that mimic transcription 159 and 174 bp into a nucleosome, a TspRI cleavage site placed either 69 or 84 bp downstream of the Widom 601 dyad. PCR reactions, purifications and TspRI digestions were performed as described above. Cleaved DNA products purified on 6% denaturing PrepCell PAGE (12% 19:1 acrylamide:bis-acrylamide, 8 M urea, 1x TBE running at 4 °C and 8 W in 0.5x TBE buffer) and subsequently concentrated by ethanol precipitation. Primers to generate the mismatch bubble (Non template 5’ –/5Phos/GCGGCCCTTGTGTTCAGGAGCCAGCAGGGAGCTGGGAGC, Template 5’ GCTCCCAGCTCCCTGCTGGCTCCGAGTGGGTTCTGCCGCGACAGTGAT) were mixed and annealed by slow cooling from 95 °C. Annealed primers were ligated to TspRI cleaved Widom 601 sequences using T4 ligase as per manufactures instructions.

DNA for tetranucleosomal arrays were generated from plasmid DNA similar to a process described previously^81^. The plasmid DNA was extracted from XL1 Blue *E. coli* cells with GigaPrep extraction kits (Macherey-Nagel) and contained four Widom 601 sequences equally spaced by 30 bp linker DNA. The plasmid DNA was digested by EcoRV-HF at 37 °C for 10 h. The desired tetranucleosomal template DNA was purified by size-selective PEG precipitation at 800 mM NaCl and approx. 6-7% PEG-6000. The DNA was digested by TspRI at 65 °C for 10 h and purified on 6% denaturing PrepCell PAGE (6% 19:1 acrylamide:bis-acrylamide, 8 M urea, 1x TBE running at 4 °C and 8 W in 0.5x TBE buffer). The ratio between DNA and histone octamer for successful reconstitution of mononucleosome or tetranucleosomes were determined by a titration series of histone octamer to DNA. The reconstitution titration was analyzed by BanI digestion, containing a restriction site in the Widom 601 sequence, and agarose PAGE. The minimal histone octamer concentration for complete inhibition of BanI cleavage was chosen to generate tetranucleosomal constructs. Nucleosome used for cryo-EM studies were reconstituted with histone octamers containing an H3K36M point mutation. The concentration of the all nucleosomal constructs were calculated by the extinction coefficient and the absorption at 280 nm.

#### RNA extension assays with methylation

RNA extension assays were performed on mononucleosomal and tetranucleosomal constructs similarly as described^6,7,33^. The mononucleosomal DNA contained T-less cassette on the template strand to nucleosomal bp 32. The modified Widom 601 sequence was located 112 bp from the 5’ end. The tetranucleosomal DNA consisted of a 35 bp run-up sequence followed by four Widom 601 sequences spaced by 30 bp linker sequences.

Mononucleosome extension assay in Fig. 1b were performed in a final volume of 18 µL. RNA (150 nM) containing a 5’ Cy5 label and nucleosomal template (60 nM) were mixed and incubated for 10 min on ice. *S. scrofa* Pol II (150 nM) was added to the reaction and incubated for 10 min on ice. DSIF (225 nM), SPT6 (225 nM), RTF1 (225 nM), PAF1c (225 nM), P-TEFb (250 nM), IWS1 (0.450 nM), FACT (300 nM), 3’d ATP (0.5 mM), SETD2 (6 µM), compensation buffer, water were added to the reaction. The reactions were incubated for 15 min at 30 °C. Transcription was started with the addition of TFIIS (90 nM), CTP, GTP, UTP, ATP (300 µM each) and S-Adenosyl methionine (SAM) (10 µM). Nucleotides or ATP were omitted from reactions without transcription or to stall the Pol II. Buffer composition in the final reaction was 90 mM NaCl, 20 mM Na·HEPES pH 7.4 (@ 20 °C), 4 mM MgCl2, 4% (v/v) glycerol, and 1 mM DTT, and 10 µM ZnCl2. Reactions were allowed to proceed for 10 min at 30 °C. To monitor RNA extension, five microliters of reactions were taken and quenched with 2x stop buffer (6.4 M urea, 50 mM EDTA pH 8.0, 2x TBE buffer). Proteinase K (40 µg) (New England Biolabs) was added to the quenched sample and incubated for 30 min at 37 °C. Samples were denatured at 95 °C for 10 min and separated by denaturing PAGE (12% acrylamide:bis-acrylamide 19:1, 8 M urea, 1x TBE buffer, ran at 300 V in 0.5x TBE buffer for approx. 40 min). To monitor H3K36me3, 12 µL of the reaction was mixed with 6 µL 4x LDS dye (Thermo Scientific). Samples were denatured for 2 minutes at 95 °C and 12 µL were subjected to denaturing gradient gel electrophoresis (NuPAGE™ 12 %, Bis-Tris gel.) and subsequently transferred onto nitrocellulose membranes (GE Healthcare Life Sciences) for Western blotting against H3K36me3, that was carried out following standard protocols.

Tetranucleosome extension assays in Fig. 3c were performed in a final volume of 15 µL. RNA (150 nM) containing a 5’ Cy5 label and nucleosomal template (60 nM) were mixed and incubated for 10 min on ice. *S. scrofa* Pol II (150 nM) was added to the reaction and incubated for 10 min on ice. DSIF (225 nM), SPT6 (225 nM), RTF1 (225 nM), PAF1c (225 nM), P-TEFb (250 nM), IWS1 (0.450 nM), FACT (300 nM), ATP (1 mM), SETD2 (0.6 µM), compensation buffer, water were added to the reaction. The reactions were incubated for 15 min at 30 °C. Transcription was started with the addition of TFIIS (90 nM), CTP, GTP, UTP (300 µM each), and SAM (4 µM). Buffer composition in the final reaction was 90 mM NaCl, 20 mM Na·HEPES pH 7.4 (@ 20 °C), 4 mM MgCl2, 4% (v/v) glycerol, 1 mM DTT, and 10 µM ZnCl2. Reactions were allowed to proceed for 30 min at 30 °C. Samples from tetranulceosomal reactions were treated with DNaseI due to the longer DNA and RNA. Five microliters from tetranucleosomal reactions were mixed with 0.6 µL 10x CaCl2 buffer (100 mM Na·HEPES pH 7.4 (@ 20 °C), 50 mM CaCl2, 25 mM MgCl2) and proteinase K (40 µg). The samples were incubated for 20 min at 55 °C, cooled down to 37 °C, before 1.5 U DNaseI (RNase-free, Thermo Scientific™) were added. The samples were incubated at 37 °C for 20 min, then quenched with 2x stop buffer, denatured for 10 min at 95 °C and separated by 6% denaturing PAGE gels (6% acrylamide:bis-acrylamide 19:1, 8 M urea, 1x TBE buffer, ran at 300 V in 0.5x TBE buffer for approx. 30 min).

#### Filter binding assays

For quantification of *in vitro* methylation in the tetranucleosome extension assays, experiments were performed as outlined above, substituting cold SAM with 4 μM 3H SAM (Perkin Elmer) 10 μL of the reaction were spotted on nitrate cellulose membranes and washed with 3 mL of 20 mM HEPES (pH 7.4) under vacuum. Membranes were dissolved and methylated products quantified by liquid scintillation. Data was normalized to the presence or absence of SETD2. Reactions were performed in triplicates. Data displayed as box plots, each point reflects one replicate (N=3), depicted as mean ± s.d.

#### Sample preparation for cryo-EM analysis

##### State 1 complex

The methylation competent elongation complex with an upstream transferred hexasome was assembled in a final buffer containing 70 mM NaCl, 20 mM Na·HEPES pH 7.4 (@ 20m °C), 3 mM MgCl2, 1 mM DTT, and 4% (v/v) glycerol. The reaction volume was 300 µL. The complex formation was performed similarly to the RNA extension assays. 5’ Cy5 labeled RNA (960 nM) and nucleosomal template (480 nM) was mixed and incubated on ice for 10 min. *S. scrofa* Pol II (400 nM) was added and the reaction was incubated on ice for 10 min. DSIF (800 nM), SPT6 (800 nM), PAF1c (800 nM), RTF1 (800 nM), P-TEFb (400 nM), AKT3 (400 nM), IWS1 (2.4 µM), SETD2 (1.6 µM), 3’dATP (0.5 mM) (Jena Bioscience), GTP (0.1 mM), UTP (0.1 mM), compensation buffer, water, and FACT (800 nM) were added to the reaction and incubated for 15 min at 30 °C. Transcription elongation was started with the addition of TFIIS (240 nM) and CTP (0.1 mM) and was allowed to continue for 30 min at 30°C to the end of the T-less cassette of the template strand at nucleosomal bp 64. The reaction was applied onto a 2 mL glycerol grafix gradient containing 10-30% (v/v) glycerol, 65 mM NaCl, 20 mM Na·HEPES pH 7.4 (@ 20 °C), 3 mM MgCl2, 1 mM DTT, and 0.0175% (v/v) glutaraldhyde in the heavy solution. The gradient was spun in a TLS-55 swinging rotor (Beckman Coulter) for 3 h at 55,000 xg and 4 °C. After the centrifugation, the gradient was fractionated in 100 µL steps from the top and quenched with 8 mM aspartate and 10 mM lysine for 10 min on ice. Fractions were analyzed using 12% denaturing PAGE (as described in RNA extension) and 3-12% nativePAGE™ (Invitrogen™). Fractions containing the complex (Extended Data Fig. 1) were quenched with 16 mM aspartate and 4 mM lysine and dialyzed for 3 h in 50 mM NaCl, 20 mM Na·HEPES, pH 7.4 (@ 20 °C), 20 mM Tris-Cl, pH 7.5 (@ 4°C), and 1 mM DTT.

For grid preparation, R2/2 UltrAuFoil grids (Quantifoil) were glow-discharged for 100 s. DDM was added to the dialyzed sample to a final concentration of 0.0025%. 3 µL sample was applied to both sides of the grids and incubated for approx. 5 s at 4 °C and 100% humidity. The grids were blotted with a blot force of 5 for 3 s before plunging into liquid ethan using a Vitrobot Mark IV (Thermo Fisher). Grids were clipped and stored in liquid nitrogen until cryo-EM analysis.

##### State 2 complex

The methylation competent elongation complex with SETD2 bound to the proximal H3 tail of an upstream nucleosome was assembled in a final buffer containing 80 mM NaCl, 20 mM Na·HEPES pH 7.4 (@ 20 °C), 3 mM MgCl2, 1 mM DTT, and 4% (v/v) glycerol. The reaction volume was 90 µL. 5’ FAM labeled RNA (1.67 µM)(/56-FAM/rUrUrArArGrGrArArUrUrArArGrUrCrGrUrGrCrGrUrCrUrArArUrArArCrCrGrGrAr GrArGrGrGrArArCrCrCrArCrU) and the ligated (+159 bp) nucleosomal template (0.86 µM) was mixed with SETD2 (8.6 µM) and SAM (0.1 mM) on ice for 10 min. *S. scrofa* Pol II (1 µM) was added and the reaction was incubated for a further 10 min on ice. DSIF (1.5 µM), SPT6 (1.5 µM), IWS1 (1.5 µM), RTF1 (2 µM), PAF1c (1.5 µM), P-TEFb (0.4 µM), ATP (0.5 mM) compensation buffer, water were added and sample incubated at 30 °C for 15 minutes. After centrifugation, the complex was purified by gel filtration using a Superose 6 Increase (3.2/300) pre-equilibrated with 50 mM NaCl, 20 mM Na·HEPES pH 7.4 (@ 20 °C), 3 mM MgCl2, 1 mM DTT, and 4% glycerol, in the presence of FACT (Extended Data Fig. 11a). A single peak fraction containing all components was with crosslinked with 0.1% w/v glutaraldehyde on ice for 10 minutes before quenching with 16 mM aspartate and 4 mM lysine.

For grid preparation, R2/2 UltrAuFoil grids (Quantifoil) were glow-discharged for 100 s. 2.5 µL sample was applied to both sides of the grids and incubated for approx. 15 s at 4°C and 100% humidity. The grids were blotted with a blot force of 5 for 3 s before plunging into liquid ethan using a Vitrobot Mark IV (Thermo Fisher).

##### State 3 complex

The methylation competent elongation complex with SETD2 bound to the distal H3 tail of an upstream nucleosome was assembled on ligated a nucleosomal template (+174 bp) and the complex purified and frozen in an identical manner to the proximal H3 bound complex.

#### Cryo-EM analysis and image processing

For State 1, cryo-EM data were collected under near-identical conditions on two separate grids during different collection periods. Data was acquired at a nominal magnification of 81,000×, corresponding to a calibrated pixel size of 1.05 Å/pixel, using a K3 direct electron detector (Gatan) on a Titan Krios transmission electron microscope (Thermo Fisher Scientific) operated at 300 kV. Images were collected in EFTEM mode using a Quantum LS energy filter with a slit width of 20 eV. Images were collected in electron counting mode with an applied defocus range of –0.5 to –2.0 μm. The SerialEM^82^ software was used for automated data acquisition. All pre-processing of the collected movies (motion correction, dose weighting, CTF estimation and particle picking) were performed using Warp^83^.

For dataset 1, we collected 45,705 micrographs with a dose rate of 14.45 e^−^/pixel/s for 3.05 s resulting in a total dose of 39.98 e^−^/Å^2^ that was fractionated into 40 movie frames. Micrographs with bad CTF fits in Warp were excluded from further processing. We extracted 5,835,867 picked particles with a box size of 512 pixels and binned 5x to a pixel size of 5.25 Å/pixel using RELION 3.1^84^. These particles were subjected to interactive rounds of 2D classification, and heterogenous refinement in CryoSPARC^85^ using initial models generated by ab initio reconstruction. Classes with good particles for RNA polymerase were re-extracted in RELION^84^ (binned 2x, pixel size 2.1 Å/pixel) and further cleaned using 2D and 3D classification. Finally, particles were re-extracted without binning and focused refinement, with a mask around RNA polymerase, was performed. These particles (432,698) were subjected to iterative rounds of CTF refinement and Bayesian polishing resulting in 3.62Å reconstruction, encompassing the activated elongation complex and additional upstream density. For dataset 2, we collected 55,723 micrographs with a dose rate of 15.83 e^−^/pixel/s for 2.82 s resulting in a total dose of 40.49 e^−^/Å2 that was fractionated into 40 movie frames. Micrograph curation, pre-processing and initial particle cleanup was performed as described in dataset1. After CTF refinement and polishing, 804,464 particles contributed to a 2.92 Å reconstruction and was merged with dataset 1. Polishing artifacts were present in both datasets were filtered based on X,Y coordinate on the micrographs.

To determine the composition of the upstream density, we applied a generous soft mask around the upstream half of Pol II and subtracted the signal outside of this region using RELION 3.1 ^67,84,86^. Subsequent 3D classification and refinement identified a subset of particles that produced of 3.73 Å resolution reconstruction (MAP 1) from which we could observe a FACT bound hexasome, immediately upstream of the polymerase. To improve the resolution, we repeated the signal subtraction, using a tighter mask around the FACT-hexasome complex, and repeated 3D classification and refinement. The resulting reconstruction (MAP 3) yielded improved local resolution and an overall reconstruction at 3.7 Å resolution. To obtain an overall reconstruction of RNA polymerase II and the FACT hexasome complex, particles from MAP 2 (after reversion of the signal subtraction) were back-projected using the angles obtained from the refinement of MAP 1 in RELION^84^. This produced the best consensus map of both the polymerase and FACT-hexasome complex at 6.4 Å resolution. To resolve the flexibility in SPT6, signal was subtracted outside a soft mask around SPT6. Subsequent 3D classification and refinement identified a two subset of particles, one containing canonical SPT6^10^ and a second subset in which SPT6 rotated upwards (MAP 4). Reversion of the signal subtraction of both subsets and subsequent global refinement resulted in one map with a stable FACT-bound hexasome (for the rotated SPT6 subset) and the second map without density for FACT or a hexasome. ∼40% of particles from MAP 4 overlaps with MAP 2, whilst only ∼2.5% of particles from the canonical SPT6 position overlap with MAP 4.

For State 2, cryo-EM data were collected under near-identical conditions to above, including nominal magnification, pixel size, detector, energy filter and defocus range. All pre-processing of the collected movies (motion correction, dose weighting, CTF estimation and particle picking) were performed using Warp^83^. We collected 77,138 micrographs with a dose rate of 16.39 e^−^/pixel/s for 2.43 s resulting in a total dose of 39.83 e^−^/Å^2^ that was fractionated into 40 movie frames. Micrographs with bad CTF fits in Warp were excluded from further processing. We extracted 4,613,946 picked particles with a box size of 512 pixels and binned 4x to a pixel size of 4.2 Å/pixel using RELION 3.1^67,84,86^. These particles were subjected to interactive rounds of 2D classification, and heterogenous refinement in CryoSPARC^85^ using initial models generated by ab initio reconstruction. Classes with good particles for RNA polymerase were re-extracted in RELION (binned 2x, pixel size 2.1 Å/pixel) and further cleaned using 2D and 3D classification and decreasing spherical masks around Pol II. Finally, particles were re-extracted without binning and focused refinement These particles (∼750,00) were subjected to iterative rounds of CTF refinement and Bayesian polishing resulting in 2.63 Å reconstruction, encompassing the activated and additional upstream density.

To determine the composition of the upstream density, we applied a generous soft mask around the upstream density and subtracted the signal outside of this region using RELION 3.1^67,84,86^. Due to severe heterogeneity, initial angular assignments required 2D classification (performed in CryoSPARC) and addition rounds of heterogenous refinement. Well aligning particles of SETD2 bound to the nucleosome were imported back into RELION 3.1^67,86^ for additional focused classification (Extended Data Fig. 10c) to the remove ∼20,000 particles that contained two SETD2 molecules bound. Remaining particles resulted in an overall reconstruction of SETD2 bound to the nucleosome at 4.25 Å from approximately 80,000 particles (MAP 1). These particles were selected from the consensus refinement stack, thereby recovering angles Pol II refinement angles. Multibody refinement performed in RELION 3.1^67^ was used to the determine principal components of nucleosome motion, relative to Pol II. Given the unimodal distribution, a subset of particles was selected based on the eigenvalue for component 1 (+/-10) and re-extracted to centre the particles between Pol II and the nucleosome. Final refinement of the centered particles was performed using a spherical mask of 250 Å diameter (MAP 3). For classification of SPT6, an initial soft mask around SPT6 was used for subtraction. Focused 3D classification with local angular sampling was used to select a subset of well aligning particles. Refinement steps that utilized Blush in RELION 5^87^ are indicated in Extended Data Fig. 10c. A second focused classification, without image shift alignments, was performed using a tight spherical mask around the DLD of SPT6. Subsequent refinement of particles containing SETD2 resulted in a 3.82 Å reconstruction (MAP 2). 18.5% of particles in MAP 1 overlap with MAP 2, suggesting a subset of the SETD2 particles remain tethered to SPT6 whilst simultaneously binding the nucleosome.

For State 3, cryo-EM data was collected in an identical fashion to State 2. All pre-processing of the collected movies (motion correction, dose weighting, CTF estimation and particle picking) were performed using Warp ^83^. We collected 74,840 micrographs with a dose rate of 16.40 e^−^/pixel/s for 2.44 s resulting in a total dose of 40.01 e^−^/Å^2^ that was fractionated into 40 movie frames. Micrographs with bad CTF fits in Warp were excluded from further processing. We extracted 9,589,557 picked particles with a box size of 512 pixels and binned 4x to a pixel size of 4.2 Å/pixel using RELION 3.1^67,84,86^. Data processing was carried out in a similar fashion to State 2 and as outlined in Extended Data Fig. 14.

#### Model building and refinement

To build a model for State 1 into MAP 2, Alphafold2 models of human SSRP1 and SPT16 were placed within the cryo-EM density using ISOLDE flexible fitting^88^. To guild fitting, PDB 7XTI^27^ was used as a reference to restrain the Alphafold2 models to. The nucleosome (PDB 2CV5)^89^ was rigid body docked into the density, using ChimeraX ^90,91^ and the proximal H2A– H2B dimer and DNA removed. Due to the anisotropic nature of the density the sequence register of the RNA polymerase II active site (in MAP 3) could not be determined. Therefore, the DNA was modelled based on the designed stall site located 64 bp into the nucleosome. ISOLDE ^88^ was used to fit the parts of SPT4 and SPT6, present in the reconstruction, from their respective models in 6TED. To model the FACT “fastener”, we used ColabFold AlphaFold2 w/ MMseqs2^58^ to generate a model of the SPT16 (residues 457-930) and RTF (residues 266-315) that was subsequently used as a guide to place the FACT “fastener” into the observed density. Similarly, ColabFold^58^ was used to model a hexasome bound by SPT16 that assisted in the modelling of the SPT16 CTD. For building Pol II into MAP 3, the MAP 2 model was merged with PDB 6TED^10^ and protein chains outside of the map density were flexibly fitted using ISOLDE^88^. MAP 4 was used to correctly position SPT6 and build the SPT6-SETD2 interface guided by the ColabFold prediction. The Alphafold2 model of the SETD2 AID was rigid body docked into MAP 1 and flexible regions removed. Reference model restraints were applied to prevent secondary structure deterioration of the model in regions of lower local resolution. For the model of MAP 1, regions of the MAP 3 model that were outside of the subtracted reconstruction were removed and the model was adjusted to the density map in ISOLDE^88^ and COOT^92^.

To model the nucleosome in State 2, PDB 7EA8^57^ was rigid body fitted into State 2 MAP 1. Histone residues where mutated to match sequences used. For SPT6, the Alphafold2 model was fitted to PDB 6TED^10^ and adjusted with ISOLDE. Residues outside the available Cryo-EM density for State 2 MAP2 were removed. The complete model of State 2 was built by fitting PDB 6TED^10^ and the models for State 2 MAP 1 and 2 into MAP 3 and B-DNA modelled between the active site of Pol II and the unwrapped nucleosome using COOT^92^. Given the flexible nature, the AID was not modelled in State 2. The CTD of SPT16 was model based on PDB 8I17^66^. For figures, the AlphaFold2 model of the AID was rigid body docked into MAP 1 Cryo-EM density in ChimeraX^90,91^. A similar approach was taken to model State 3.

All models were refined the using phenix.real_space_refine tool in the PHENIX package^93,94^.

#### Crosslinking mass spectrometry

For crosslinking mass spectrometry, the activated elongation complex transcribed to nucleosomal base pair 64 with an upstream FACT-bound hexasome was assembled similarly to the formation for cryo-EM. The differences of the complex assembly compared to the cryo-EM sample preparation are listed below. The final volume of the assembly was 600 µL with proteins and nucleic acids scaled up accordingly. To stall the polymerase at the end of the T-less cassette, 1 mM 3’dATP (Jena Bioscience) was used.

The complex assembly was applied onto a 4 mL glycerol gradient containing 15-45% (v/v) glycerol, 65 mM NaCl, 20 mM Na·HEPES pH 7.4 (@ 20 °C), 3 mM MgCl2, and 1 mM DTT. The gradient was spun in a SW55 Ti swinging rotor (Beckman Coulter) for 16 h at 55,000 rpm and 4 °C. Post centrifugation, the gradient was fractionated in 200 µL steps from the top. Fractions containing the assembled complex were pooled and crosslinked with 2 mM bis(sulfosuccinimidyl)suberate (BS3) on ice before being quenched by 100 mM Tris-HCl pH 7.4 (@ 20 °C). The complexes were pelleted by ultracentrifugation in an S150AT rotor (Thermo Fisher Scientific). The pellet was solubilized in 50 mM ammonium bicarbonate (pH 8.0) supplemented with 4M urea, reduced with 5 mM DTT and alkylated with 17 mM iodoacetamide. The sample was diluted with 50 mM ammonium bicarbonate to reduce urea concentration to 1M and digested with trypsin (Promega) in a 1:20 enzyme-to-protein ratio (w/w) at 37 °C overnight. Peptides were reverse-phase extracted using SepPak Vac tC18 1cc/50mg (Waters) and eluted with 50% acetonitrile (ACN) / 0.1% trifluoroacetic acid (TFA). The eluate was dried in a vacuum concentrator (Eppendorf). Dried peptides derived from 33 pmol of the complex were dissolved in 35 µl of 2% ACN / 20 mM ammonium hydroxide and separated by reverse phase HPLC at basic pH using an xBridge C18 3.5µm 1×150mm column (Waters) at a flow rate of 60 µl/min at 24°C. Buffers A and B for mobile phase were 20 mM ammonium hydroxide, pH 10, and 80% ACN / 20 mM ammonium hydroxide, pH 10, respectively. Peptides were bound to a column pre-equilibrated with 5% buffer B and eluted over 64 min using the following gradient: 5%B (min 0-4), 5-8%B (min 5-7), 8-36%B (min 8-41), 36-45%B (min 42-49), 45-95%B (min 50), 95%B (min 51-55), 95-5%B (min 56-57), 5%B (min 58-63). Fractions of 60 µl were collected. Peptides eluted between minute 3 and 55 were vacuum dried and dissolved in 5% ACN / 0.1% TFA for subsequent uHPLC-ESI-MS/MS analysis. The fractions were injected into a Dionex UltiMate 3000 uHPLC system (Thermo Scientific) coupled to a Thermo Orbitrap Exploris mass spectrometer and measured twice with a 60 and twice with a 90 min method. For uHPLC, a C18 PepMAP 100 trap column (0.3 x 5 mm, 5 μm, Thermo Scientific) and a custom 30 cm C18 main column (75 µm inner diameter packed with ReproSil-Pur 120 C18-AQ beads, 3 µm pore size, Dr. Maisch GmbH) was used. Mobile phase was formed using buffers A (0.1% formic acid) and B (80% ACN / 0.08% formic acid). Peptide separation was achieved by applying a linear gradient of 11-45%B (min 3 to 42 in a 60 min method or min 3 to 72 in a 90 min method) followed by 45-52%B (min 43 to 47 in a 60 min method or min 73 to 78 in a 90 min method). MS settings were as follows: MS1 resolution, 120000; MS1 scan range, 350-1550 m/z; MS1 normalized AGC target, 300%; MS1 maximum injection time, auto; cycle time (Top Speed), 3 s; intensity threshold, 1E4; MS2 resolution, 60000; isolation window, 1.6 Th; normalized collision energy, 30%; MS2 AGC target, 75%; MS2 maximum injection time, 128 ms. Only precursors with a charge state of 3-8 were selected for MS2 using a dynamic exclusion of 25 s. Protein-protein crosslinks were identified by searching Thermo raw files against a custom database of 31 proteins using pLink2.3.11 software (http://pfind.org/software/pLink/) and filtered at false discovery rate (FDR) of 5% according to the recommendations of the developer^95,96^

#### Figure generation

Figures were generated using UCSF ChimeraX (version 1.6)^90,91^.

#### Quantification and statistical analysis

Quantification of signal from denaturing PAGE gels were performed using the integrated density function of ImageJ2 2.3.0. No statistical analysis were performed.

## DATA AND CODE AVAILABILITY

- The cryo-EM reconstructions and final models for the State 1-3, have been deposited in the Electron Microscopy Data Base under ID codes XXX, XXX and XXX, respectively, and in the Protein Data Bank under ID codes XXX, XXX and XXX respectively.
- This paper does not report original code.
- Any additional information required to re-analyze the data reported in this paper is available from the lead contact upon request.

